# An ancestral interaction module promotes oligomerization in divergent mitochondrial ATP synthases

**DOI:** 10.1101/2021.10.10.463820

**Authors:** Ondřej Gahura, Alexander Mühleip, Carolina Hierro-Yap, Brian Panicucci, Minal Jain, David Hollaus, Martina Slapničková, Alena Zíková, Alexey Amunts

**Affiliations:** Institute of Parasitology, Biology Centre CAS, Ceske Budejovice, Czech Republic; Science for Life Laboratory, Department of Biochemistry and Biophysics, Stockholm University, 17165 Solna, Sweden; Faculty of Science, University of South Bohemia, Ceske Budejovice, Czech Republic

## Abstract

Mitochondrial ATP synthase forms stable dimers arranged into oligomeric assemblies that generate the inner-membrane curvature essential for efficient energy conversion. Here, we report cryo-EM structures of the intact ATP synthase dimer from *Trypanosoma brucei* in ten different rotational states. The model consists of 25 subunits, including nine lineage-specific, as well as 36 lipids. The rotary mechanism is influenced by the divergent peripheral stalk, conferring a greater conformational flexibility. Proton transfer in the lumenal half-channel occurs via a chain of five ordered water molecules. The dimerization interface is formed by subunit-*g* that is critical for interactions but not for the catalytic activity. Although overall dimer architecture varies among eukaryotes, we find that subunit-*g* together with subunit-*e* form an ancestral oligomerization motif, which is shared between the trypanosomal and mammalian lineages. Therefore, our data defines the subunit-*g/e* module as a structural component determining ATP synthase oligomeric assemblies.

---

Mitochondrial ATP synthase consists of the soluble F_1_ and membrane-bound F_o_ subcomplexes, and occurs in dimers that assemble into oligomers to induce the formation of inner-membrane folds, called cristae. The cristae are the sites for oxidative phosphorylation and energy conversion in eukaryotic cells. Dissociation of ATP synthase dimers into monomers results in the loss of native cristae architecture and impairs mitochondrial function^1,2^. While cristae morphology varies substantially between organisms from different lineages, ranging from flat lamellar in opisthokonts to coiled tubular in ciliates and discoidal in euglenozoans^3^, the mitochondrial ATP synthase dimers represent a universal occurrence to maintain the membrane shape^4^.

ATP synthase dimers of variable size and architecture, classified into types I to IV have recently been resolved by high-resolution cryo-EM studies. In the structure of the type-I ATP synthase dimer from mammals, the monomers are only weakly associated^5,6^, and in yeast insertions in the membrane subunits form tighter contacts^7^. The structure of the type-II ATP synthase dimer from the alga *Polytomella* sp. showed that the dimer interface is formed by phylum-specific components^8^. The type-III ATP synthase dimer from the ciliate *Tetrahymena* thermophila is characterized by parallel rotary axes, and a substoichiometric subunit, as well as multiple lipids were identified at the dimer interface, while additional protein components that tie the monomers together are distributed between the matrix, transmembrane, and lumenal regions^9^. The structure of the type-IV ATP synthase with native lipids from *Euglena gracilis* also showed that specific protein-lipid interactions contribute to the dimerization, and that the central and peripheral stalks interact with each other directly^10^. Finally, a unique apicomplexan ATP synthase dimerizes via 11 parasite-specific components that contribute ~7000 Å^2^ buried surface area^11^, and unlike all other ATP synthases, that assemble into rows, it associates in higher oligomeric states of pentagonal pyramids in the curved apical membrane regions. Together, the available structural data suggest a diversity of oligomerization, and it remains unknown whether common elements mediating these interactions exist or whether dimerization of ATP synthase occurred independently and multiple times in evolution^4^.

The ATP synthase of *Trypanosoma brucei*, a representative of kinetoplastids and an established medically important model organism causing the sleeping sickness, is highly divergent, exemplified by the pyramid-shaped F_1_ head containing a phylum specific subunit^12,13^. The dimers are sensitive to the lack of cardiolipin^14^ and form short left-handed helical segments that extend across the membrane ridge of the discoidal cristae^15^. Uniquely among aerobic eukaryotes, the mammalian life cycle stage of *T. brucei* utilizes the reverse mode of ATP synthase, using the enzyme as a proton pump to maintain mitochondrial membrane potential at the expense of ATP^16,17^. In contrast, the insect stages of the parasite employ the ATP-producing forward mode of the enzyme^18,19^.

Given the conservation of the core subunits, the different nature of oligomerization and the ability to test structural hypotheses biochemically, we reasoned that investigation of the *T. brucei* ATP synthase structure and function would provide the missing evolutionary link to understand how the monomers interact to form physiological dimers. Here, we address this question by combining structural, functional and evolutionary analysis of the *T. brucei* ATP synthase dimer.

## Results

### Cryo-EM structure of the *T. brucei* ATP synthase

We purified ATP synthase dimers from cultured *T. brucei* procyclic trypomastigotes by affinity chromatography with a recombinant natural protein inhibitor TbIF_1_^20^, and subjected the sample to cryo-EM analysis (Extended Data Fig. 1 and 2). Using masked refinements, maps were obtained for the membrane region, the rotor, and the peripheral stalk. To describe the conformational space of the *T. brucei* ATP synthase, we resolved ten distinct rotary substates, which were refined to 3.5-4.3 Å resolution. Finally, particles with both monomers in rotational state 1 were selected, and the consensus structure of the dimer was refined to 3.2 Å resolution (Extended Data Table 1, Extended Data Figs. 2&3).

Unlike the wide-angle architecture of dimers found in animals and fungi, the *T. brucei* ATP synthase displays an angle of 60° between the two F_1_/*c*-ring subcomplexes. The model of the *T. brucei* ATP synthase includes all 25 different subunits, nine of which are lineage-specific (Fig. 1a, Supplementary Video 1, Extended Data Fig. 4). We named the subunits according to the previously proposed nomenclature^21–23^ (Extended Data Table 2). In addition, we identified and modeled 36 bound phospholipids, including 24 cardiolipins (Extended Data Fig. 5). Both detergents used during purification, n-dodecyl β-D-maltoside (β-DDM) and glyco-diosgenin (GDN) are also resolved in the periphery of the membrane region (Extended Data Fig. 6).

**Fig. 1:**
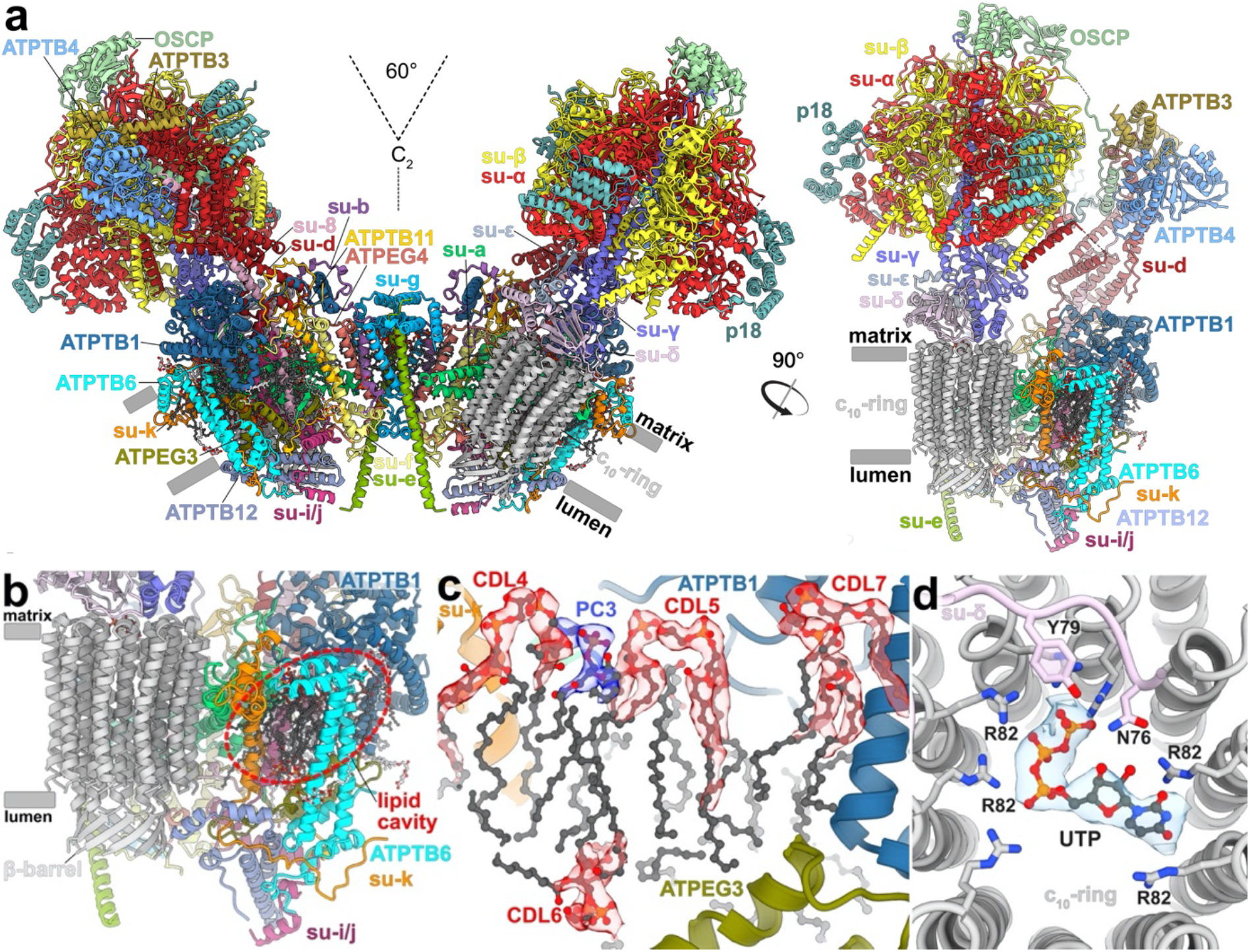
The *T. brucei* ATP synthase structure with lipids and ligands. **a**, Front and side views of the composite model with both monomers in rotational state 1. The two F_1_/*c*_10_-ring complexes, each augmented by three copies of the phylum-specific p18 subunit, are tied together at a 60°-angle. The membrane-bound F_o_ region displays a unique architecture and is composed of both conserved and phylum-specific subunits. **b**, Side view of the F_o_ region showing the lumenal interaction of the ten-stranded β-barrel of the *c*-ring (grey) with ATPTB12 (pale blue). The lipid-filled peripheral F_o_ cavity is indicated. **c**, Close-up view of the bound lipids within the peripheral F_o_ cavity with cryo-EM density shown. **d**, Top view into the decameric *c*-ring with a bound pyrimidine ribonucleoside triphosphate, assigned as UTP, although not experimentally detected. Map density shown in transparent blue, interacting residues shown.

In the catalytic region, F_1_ is augmented by three copies of subunit p18, each bound to subunit-*α*^12,13^. Our structure shows that p18 is involved in the unusual attachment of F_1_ to the peripheral stalk. The membrane region includes eight conserved F_o_ subunits (*b*, *d, f*, 8, *i*/*j*, *k*, *e*, and *g*) arranged around the central proton translocator subunit-*a*. We identified those subunits based on the structural similarity and matching topology to their yeast counterparts (Fig. 2). For subunit-*b*, a single transmembrane helix superimposes well with *b*H1 from yeast and anchors the newly identified subunit-*e* and -*g* to the F_o_ (Fig. 2a,b). In yeast and bovine ATP synthases *b*H1 and transmembrane helices of subunits-*e* and -*g* are arranged in the same way as in our structure and contribute to a characteristic wedge in the membrane domain^5^. The long helix *b*H2, which constitutes the central part of the peripheral stalk in other organisms is absent in *T. brucei* (Fig. 2c). No alternative subunit-*b*^24^ is found in our structure.

**Fig. 2:**
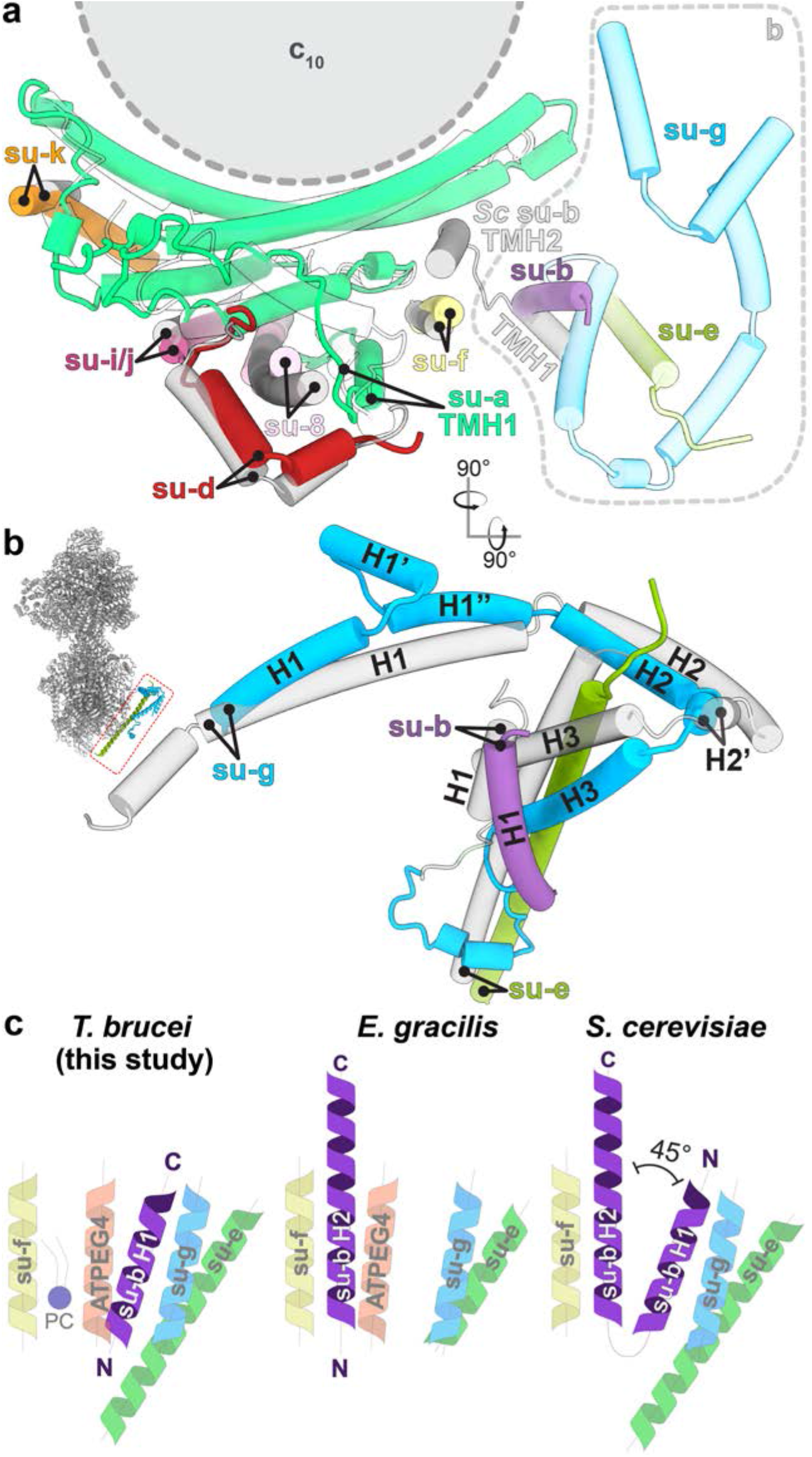
Identification of conserved F_o_ subunits. **a**, Top view of the membrane region with *T. brucei* subunits (colored) overlaid with *S. cerevisiae* structure (gray transparent). Close structural superposition and matching topology allowed the assignment of conserved subunits based on matching topology and location. **b**, Superposition of subunits-*b*, -*e* and *-g* with their *S. cerevisiae* counterparts (PDB 6B2Z) confirms their identity. **c**, Schematic representation of transmembrane helices of subunit-*b* and adjacent subunits in *T. brucei, E. gracilis* (PDB 6TDV)^10^ and *S. cerevisiae* (PDB 6B2Z)^7^ ATP synthases. PC – phosphatidylcholine.

The membrane region contains a peripheral subcomplex, formed primarily by the phylum-specific ATPTB1,6,12 and ATPEG3 (Fig. 1b). It is separated from the conserved core by a membrane-intrinsic cavity, in which nine bound cardiolipins are resolved (Fig. 1c), and the C-terminus of ATPTB12 interacts with the lumenal β-barrel of the *c*_10_-ring. The β-barrel, which has previously been reported also in the ATP synthase from *E. gracilis*^10^, extends from the *c*_10_-ring approximately 15 Å to the lumen (Fig. 1a and Extended Data Fig. 7). The cavity of the decameric *c*-ring contains density consistent with disordered lipids, as observed in other ATP synthases^5,6,7^, and in addition near the matrix side, 10 Arg66c residues coordinate a ligand density, which is consistent with a pyrimidine ribonucleoside triphosphate (Fig. 1d). We assign this density as uridine-triphosphate (UTP), due to its large requirement in the mitochondrial RNA metabolism of African trypanosomes being a substrate for post-transcriptional RNA editing^25^, and addition of poly-uridine tails to gRNAs and rRNAs^26,27^, as well as due to low abundance of cytidine triphosphate (CTP)^28^. The nucleotide base is inserted between two Arg82_c_ residues, whereas the triphosphate region is coordinated by another five Arg82_c_ residues, with Tyr79_δ_ and Asn76_δ_ providing asymmetric coordination contacts. The presence of a nucleotide inside the *c*-ring is surprising, given the recent reports of phospholipids inside the *c*-rings in mammals^5,6^ and ciliates^9^, indicating that a range of different ligands can provide structural scaffolding.

### Peripheral stalk flexibility and distinct rotational states

The trypanosomal peripheral stalk displays a markedly different architecture compared to its yeast and mammalian counterparts. In the opisthokont complexes, the peripheral stalk is organized around the long *b*H2, which extends from the membrane ~15 nm into the matrix and attaches to OSCP at the top of F_1_^5,7^. By contrast, *T. brucei* lacks the canonical *b*H2 and instead, helices 5-7 of divergent subunit-*d* and the C-terminal helix of extended subunit-*8* bind to a C-terminal extension of OSCP at the apical part of the peripheral stalk (Fig. 3a). The interaction between OSCP and subunit-*d* and *-8* is stabilized by soluble ATPTB3 and ATPTB4. The peripheral stalk is rooted to the membrane subcomplex by a transmembrane helix of subunit-*8*, wrapped on the matrix side by helices 8-11 of subunit-*d*. Apart from the canonical contacts at the top of F_1_, the peripheral stalk is attached to the F_1_ via a euglenozoa-specific C-terminal extension of OSCP, which contains a disordered linker and a terminal helix hairpin extending between the F_1_-bound p18 and subunits *-d* and -*8* of the peripheral stalk (Fig. 3a, Supplementary Videos 2,3). Another interaction of F_1_ with the peripheral stalk occurs between the stacked C-terminal helices of subunit-β and -*d* (Fig. 3b), the latter of which structurally belongs to F_1_ and is connected to the peripheral stalk via a flexible linker.

**Fig. 3:**
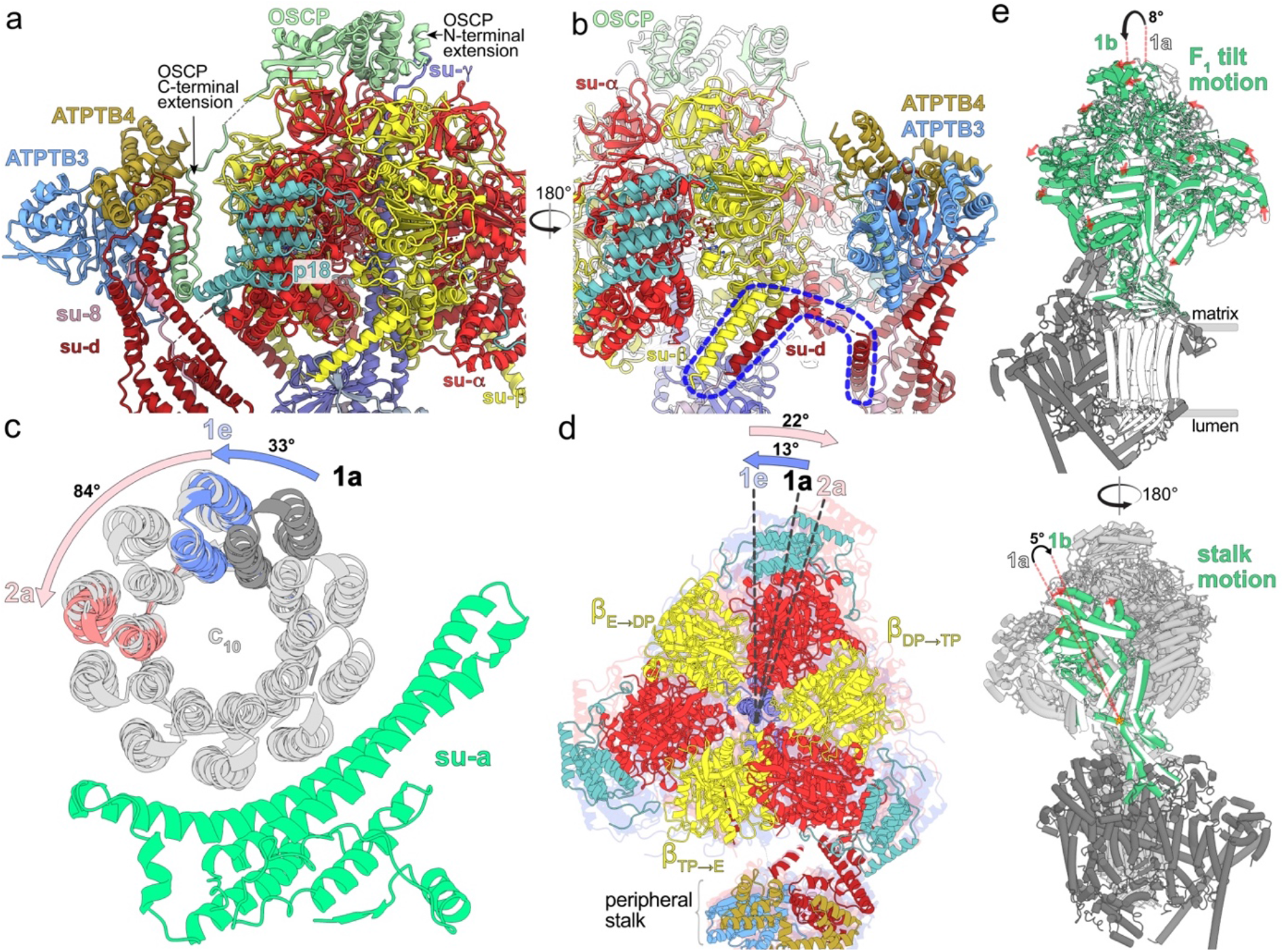
A divergent peripheral stalk allows high flexibility during rotary catalysis. **a**, N-terminal OSCP extension provides a permanent central stalk attachment, while the C-terminal extension provides a phylum-specific attachment to the divergent peripheral stalk. **b**, The C-terminal helices of subunits *-β* and -*d* provide a permanent F_1_ attachment. **c**, Substeps of the *c*-ring during transition from rotational state 1 to 2. **d**, F_1_ motion accommodating steps shown in (c). After advancing along with the rotor to state 1e, the F_1_ rotates in the opposite direction when transitioning to state 2a. **e**, Tilting motion of F_1_ and accommodating bending of the peripheral stalk.

To assess whether the unusual peripheral stalk architecture influences the rotary mechanism, we analysed 10 classes representing different rotational states. The three main states (1-3) result from three ~120° rotation steps of the rotor relatively to the static F_o_. In all classes F_1_ is in a similar conformation, corresponding to the catalytic dwell, observed previously also in the crystal structure of *T. brucei* F_1_-ATPase^13^. In accordance with the ~120° rotation of the central stalk, the conformations and nucleotide occupancy of the catalytic interfaces of the individual *aβ* dimers differ between the main states, showing ADP and ATP in the “loose” and “tight” closed conformations, respectively, and empty nucleotide binding site in the “open” conformation. We identified five (1a-1e), four (2a-2d) and one (3) classes of the respective main states. The rotor positions of the rotational states 1a, 2a and 3 are related by steps of 117°, 136° and 107°, respectively. Throughout all the identified substeps of the rotational state 1 (classes 1a to 1e) the rotor turns by ~33°, which corresponds approximately to the advancement by one subunit-*c* of the *c*_10_-ring (Fig. 3c). While rotating along with the rotor, the F_1_ headpiece lags behind, advancing by only ~13°. During the following transition from 1e to 2a, the rotor advances by ~84°, whereas the F_1_ headpiece rotates ~22° in the opposite direction (Fig. 3d). This generates a counter-directional torque between the two motors, which is consistent with a power-stroke mechanism. This counter-directional torque may occur in all three main rotational state transitions. However, it was observed only in the main state 1, because it was captured in more substeps than the remaining two states, presumably as a consequence of the symmetry mismatch between the decameric *c*-ring and the *a*_3_*β*_3_ hexamer^29^. Within the four classes of the state 2 the rotor advances by 23° and F_1_ returns close to its position observed in class 1a, where it is found also in the only observed class of the state 3. Albeit with small differences in step size, this mechanism is consistent with a previous observation in the *Polytomella* ATP synthase^8^. However, due to its large, rigid peripheral stalk, the *Polytomella* ATP synthase mainly displays rotational substeps, whereas the *Trypanosoma* F_1_ also displays a tilting motion of ~8° revealed by rotary states 1a and 1b (Fig. 3e, Supplementary Video 2). The previously reported hinge motion between the N- and C-terminal domains of OSCP^8^ is not found in our structures, instead, the conformational changes of the F_1_/*c*_10_-ring subcomplex are accommodated by a 5° bending of the apical part of the peripheral stalk. (Fig. 3e, Supplementary Videos 2,3). Together, the structural data indicate that the divergent peripheral stalk attachment confers greater conformational flexibility to the *T. brucei* ATP synthase.

### Lumenal proton half-channel is insulated by a lipid and contains ordered water molecules

The mechanism of proton translocation involves sequential protonation of E102 of subunits-*c*, rotation of the *c*_10_-ring with neutralized E102*c* exposed to the phospholipid bilayer, and release of protons on the other side of the membrane. The sites of proton binding and release are separated by the conserved R146 contributed by the horizontal helix H5 of subunit-*a* and are accessible from the cristae lumen and mitochondrial matrix by aqueous half-channels (Fig. 4a). Together, R146 and the adjacent N209 coordinate a pair of water molecules in between helices H5 and H6 (Fig. 4b). A similar coordination has been observed in the *Polytomella* ATP synthase^8^. The coordination of water likely restricts the R146 to rotamers that extend towards the *c*-ring, with which it is thought to interact.

**Fig. 4:**
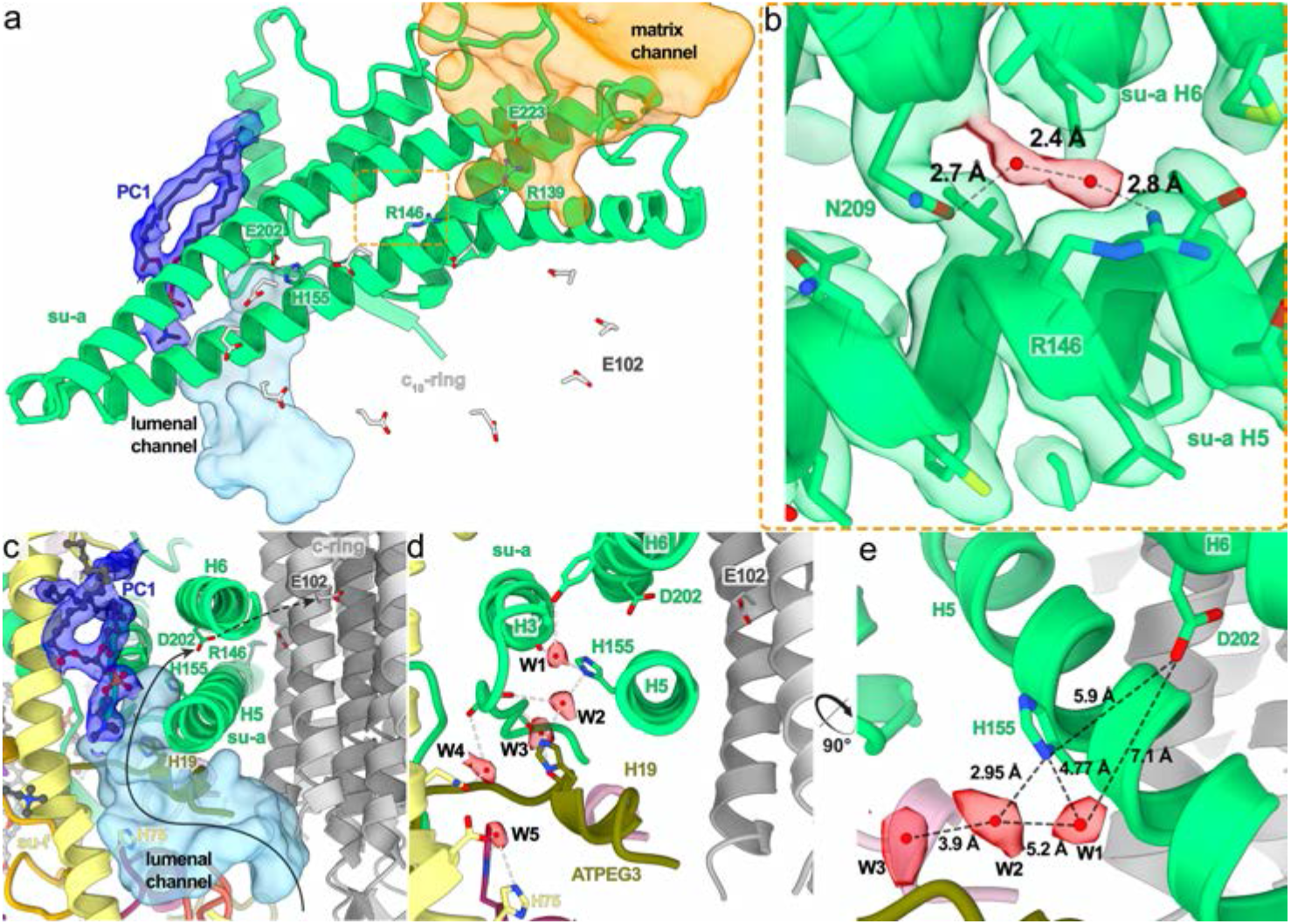
The lumenal half-channel contains ordered water molecules and is confined by an Fo-bound lipid. **a**, Subunit-*a* (green) with the matrix (orange) and lumenal (light blue) channels, and an ordered phosphatidylcholine (PC1; blue). E102 of the *c*_10_-ring shown in grey. **b**, Close-up view of the highly conserved R146_*a*_ and N209_*a*_, which coordinate two water molecules between helices H5-6_*a*_. **c**, Sideview of the lumenal channel with proton pathway (light blue) and confining phosphatidylcholine (blue). **d**, Chain of ordered water molecules in the lumenal channel. Distances between the W1-W5 (red) are 5.2, 3.9, 7.3 and 4.8 Å, respectively. **e**, The ordered waters extend to H155_*a*_, which likely mediates the transfer of protons to D202_*a*_.

In our structure, the lumenal half-channel, which displays a local resolution of 2.55 Å (Extended Data Fig. 3), is filled with a network of resolved water densities, ending in a chain of five ordered water molecules (W1-W5; Fig. 4c,d,e). The presence of ordered water molecules in the aqueous channel is consistent with a Grotthuss-type mechanism for proton transfer, which would not require long-distance diffusion of water molecules^5^. However, because some distances between the observed water molecules are too large for direct hydrogen bonding, proton transfer may involve both coordinated and disordered water molecules. The distance of 7 Å between the last resolved water (W1) and D202_*a*_, the conserved residue that is thought to transfer protons to the *c*-ring, is too long for direct proton transfer. Instead, it may occur via the adjacent H155_*a*_. Therefore, our structure resolves individual elements participating in proton transport (Fig. 4d,e).

The lumenal proton half-channel in the mammalian^5,6^ and apicomplexan^11^ ATP synthase is lined by the transmembrane part of *b*H2, which is absent in *T. brucei*. Instead, the position of *b*H2 is occupied by a fully ordered phosphatidylcholine in our structure (PC1; Fig. 4a,c). Therefore, a bound lipid replaces a proteinaceous element in the proton path.

### Subunit-*g* facilitates assembly of different ATP synthase oligomers

Despite sharing a set of conserved F_o_ subunits, the *T. brucei* ATP synthase dimer displays a markedly different dimer architecture compared to previously determined structures. First, its dimerization interface of 3,600 Å^2^ is smaller than that of the *E. gracilis* type-IV (10,000 Å^2^) and the *T. thermophila* type-III ATP synthases (16,000 Å^2^). Second, unlike mammalian and fungal ATP synthase, in which the peripheral stalks extend in the plane defined by the two rotary axes, in our structure the monomers are rotated such that the peripheral stalks are offset laterally on the opposite sides of the plane. Due to the rotated monomers, this architecture is associated with a specific dimerization interface, where two subunit-*g* copies interact homotypically on the C_2_ symmetry axis (Fig. 5a, Supplementary Video 1). Both copies of H1-2_g_ extend horizontally along the matrix side of the membrane, clamping against each other (Fig. 5c,e). This facilitates formation of contacts between an associated transmembrane helix of subunit-*e* with the neighbouring monomer via subunit-*a’* in the membrane, and *-f’* in the lumen, thereby further contributing to the interface (Fig. 5b). Thus, the ATP synthase dimer is assembled via the subunit-*e*/*g* module. The C-terminal part of the subunit-*e* helix extends into the lumen, towards the ten-stranded β-barrel of the *c*-ring (Extended Data Fig. 7a). The terminal 23 residues are disordered with poorly resolved density connecting to the detergent plug of the *c*-ring β-barrel (Extended Data Fig. 7b). This resembles the lumenal C-terminus of subunit-*e* in the bovine structure^5^, indicating a conserved interaction with the *c*-ring. In mammals, a mechanism, in which retraction of subunit-*e* upon calcium exposure pulls out the lipid plug and induces disassembly of the c-ring, which triggers permeability transition pore (PTP) opening, has been proposed^6^.

**Fig. 5:**
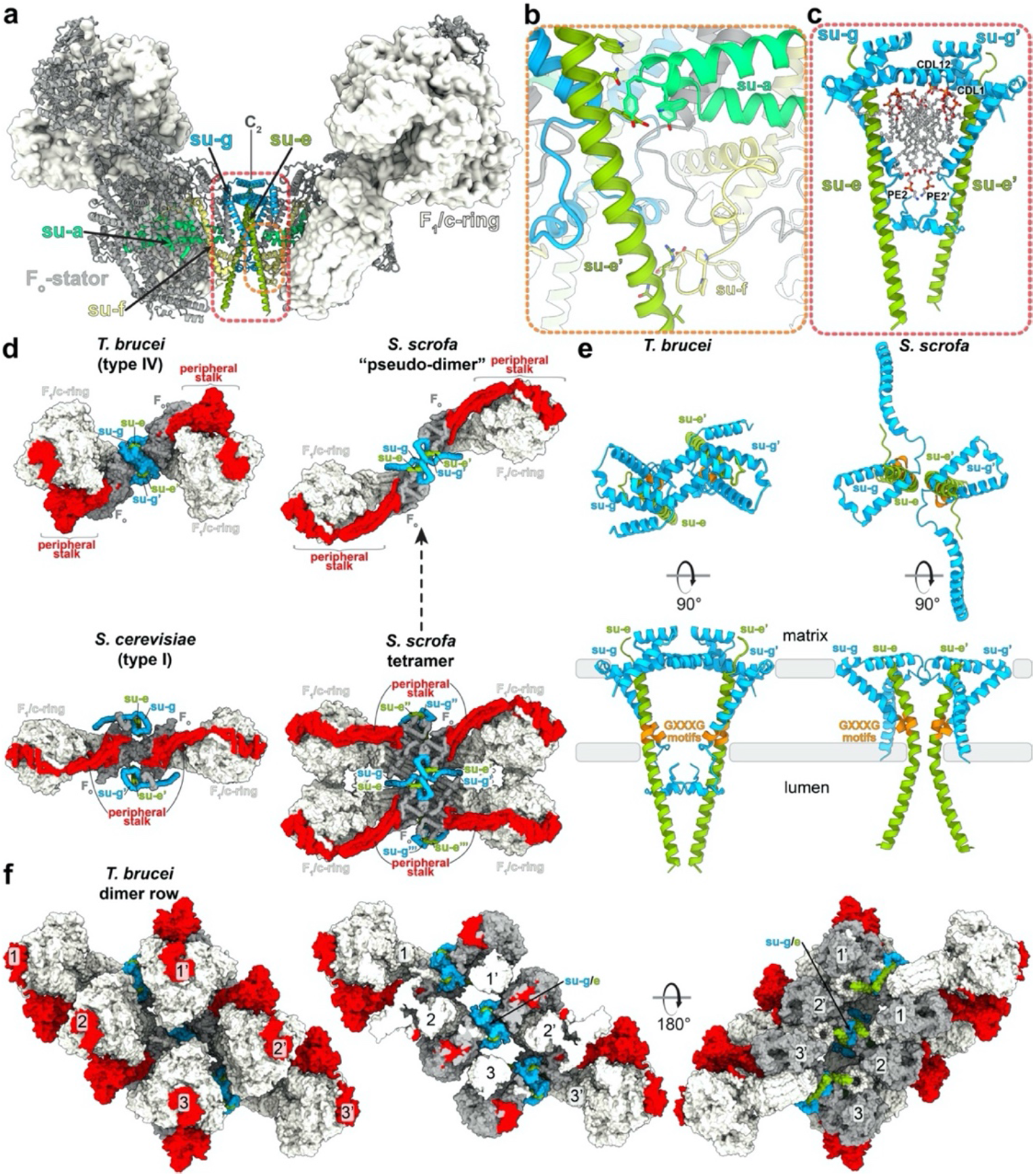
The homotypic dimerization motif of subunit-*g* generates a conserved oligomerization module. **a**, Side view with dimerising subunits colored. **b,c**, The dimer interface is constituted by (b) subunit-*e’* contacting subunit-*a* in the membrane and subunit-*f* in the lumen, (c) subunits *e* and *g* from both monomers forming a subcomplex with bound lipids. **d**, Subunit-*g* and -*e* form a dimerization motif in the trypanosomal (type-IV) ATP synthase dimer (this study), the same structural element forms the oligomerization motif in the porcine ATP synthase tetramer. The structural similarity of the pseudo-dimer (i.e., two diagonal monomers from adjacent dimers) in the porcine structure with the trypanosomal dimer suggests that type I and IV ATP synthase dimers have evolved through divergence from a common ancestor. **e**, The dimeric subunit-*e/g* structures are conserved in pig (PDB 6ZNA) and *T. brucei* (this work) and contain a conserved GXXXG motif (orange) mediating interaction of transmembrane helices. **f**, Models of the ATP synthase dimers fitted into subtomogram averages of short oligomers^15^: matrix view, left; cut-through, middle, lumenal view, right (EMD-3560).

The *e/g* module is held together by four bound cardiolipins in the matrix leaflet, anchoring it to the remaining F_o_ region (Fig. 5c). The head groups of the lipids are coordinated by polar and charged residues with their acyl chains filling a central cavity in the membrane region at the dimer interface (Fig 5c, Extended Data Fig. 5f). Cardiolipin binding has previously been reported to be obligatory for dimerization in secondary transporters^30^ and the depletion of cardiolipin synthase resulted in reduced levels of ATP synthase in bloodstream trypanosomes^14^.

Interestingly, for yeasts, early blue native gel electrophoresis^31^ and subtomogram averaging studies^2^ suggested subunit-*g* as potentially dimer-mediating, however the *e/g* modules are located laterally opposed on either side of the dimer long axis, in the periphery of the complex, ~8.5 nm apart from each other. Because the *e/g* modules do not interact directly within the yeast ATP synthase dimer, they have been proposed to serve as membrane-bending elements, whereas the major dimer contacts are formed by subunit-*a* and -*i/j*^7^. In mammals, the *e/g* module occupies the same position as in yeasts, forming the interaction between two diagonal monomers in a tetramer^5,6,32^, as well as between parallel dimers^33^. The comparison with our structure shows that the overall organization of the intra-dimeric trypanosomal and inter-dimeric mammalian *e/g* module is structurally similar (Fig. 5d). Furthermore, kinetoplastid parasites and mammals share conserved GXXXG motifs in subunit-*e*^34^ and -*g* (Extended Data Fig. 8), which allow close interaction of their transmembrane helices (Fig. 5e), providing further evidence for subunit homology. However, while the mammalian ATP synthase dimers are arranged perpendicularly to the long axis of their rows along the edge of cristae^35^, the *T. brucei* dimers on the rims of discoidal cristae are inclined ~45° to the row axis^15^. Therefore, the *e/g* module occupies equivalent positions in the rows of both evolutionary distant groups (Fig. 5f and reference 33).

### Subunit-*g* retains the dimer but is not essential for the catalytic monomer

To validate structural insights, we knocked down each individual F_o_ subunit by inducible RNA interference (RNAi). All target mRNAs dropped to 5-20 % of their original levels after two and four days of induction (Fig. 6a and Extended Data Fig. 9a). Western blot analysis of whole-cell lysates resolved by denaturing electrophoresis revealed decreased levels of F_o_ subunits ATPB1 and *-d* suggesting that the integrity of the F_o_ moiety depends on the presence of other F_o_ subunits (Fig. 6c,d). Immunoblotting of mitochondrial complexes resolved by blue native polyacrylamide gel electrophoresis (BN-PAGE) with antibodies against F_1_ and F_o_ subunits revealed a strong decrease or nearly complete loss of dimeric and monomeric forms of ATP synthases four days after induction of RNAi of most subunits (*b*, *e*, *f*, *i*/*j*, *k*, 8, ATPTB3, ATPTB4, ATPTB6, ATPTB11, ATPTB12, ATPEG3 and ATPEG4), documenting an increased instability of the enzyme or defects in its assembly. Simultaneous accumulation in F_1_-ATPase, as observed by BN-PAGE, demonstrated that the catalytic moiety remains intact after the disruption of the peripheral stalk or the membrane subcomplex (Fig. 6b,c,d and Extended Data Fig. 9b).

**Fig. 6:**
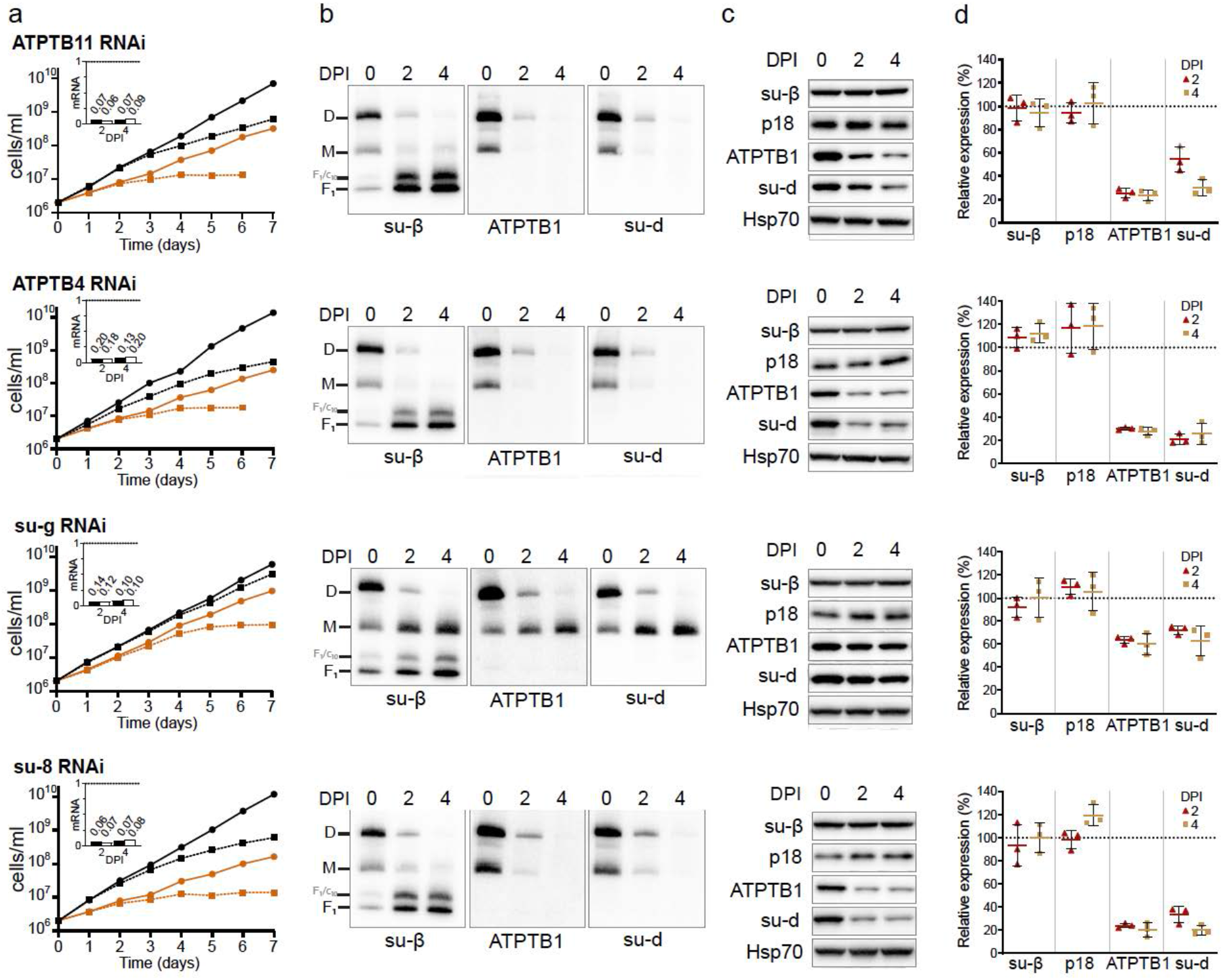
RNAi knockdown of subunit-*g* results in monomerization of ATP synthase. **a**, Growth curves of non-induced (solid lines) and tetracycline-induced (dashed lines) RNAi cell lines grown in the presence (black) or absence (brown) of glucose. The insets show relative levels of the respective target mRNA at indicated days post-induction (DPI) normalized to the levels of 18S rRNA (black bars) or β-tubulin (white bars). **b**, Immunoblots of mitochondrial lysates from indicated RNAi cell lines resolved by BN-PAGE probed with antibodies against indicated ATP synthase subunits. **c**, Representative immunoblots of whole cell lysates from indicated RNAi cell lines probed with indicated antibodies. **d**, Quantification of three replicates of immunoblots in (c). Values were normalized to the signal of the loading control Hsp70 and to non-induced cells. Plots show means with standard deviations (SD).

In contrast to the other targeted F_o_ subunits, the downregulation of subunit-*g* with RNAi resulted in a specific loss of dimeric complexes with concomitant accumulation of monomers (Fig. 6b), indicating that it is required for dimerization, but not for the assembly and stability of the monomeric F_1_F_o_ ATP synthase units. Transmission electron microscopy of thin cell sections revealed that the ATP synthase monomerization in the subunit-*g*^RNAi^ cell line had the same effect on mitochondrial ultrastructure as nearly complete loss of monomers and dimers upon knockdown of subunit-8. Both cell lines exhibited decreased cristae counts and aberrant cristae morphology (Fig. 7a,b), including the appearance of round shapes reminiscent of structures detected upon deletion of subunit-*g* or -*e* in *Saccharomyces cerevisiae*^1^. These results indicate that monomerization prevents the trypanosomal ATP synthase from assembling into short helical rows on the rims of the discoidal cristae^15^, as has been reported for impaired oligomerization in counterparts from other eukaryotes^2,36^.

**Fig. 7:**
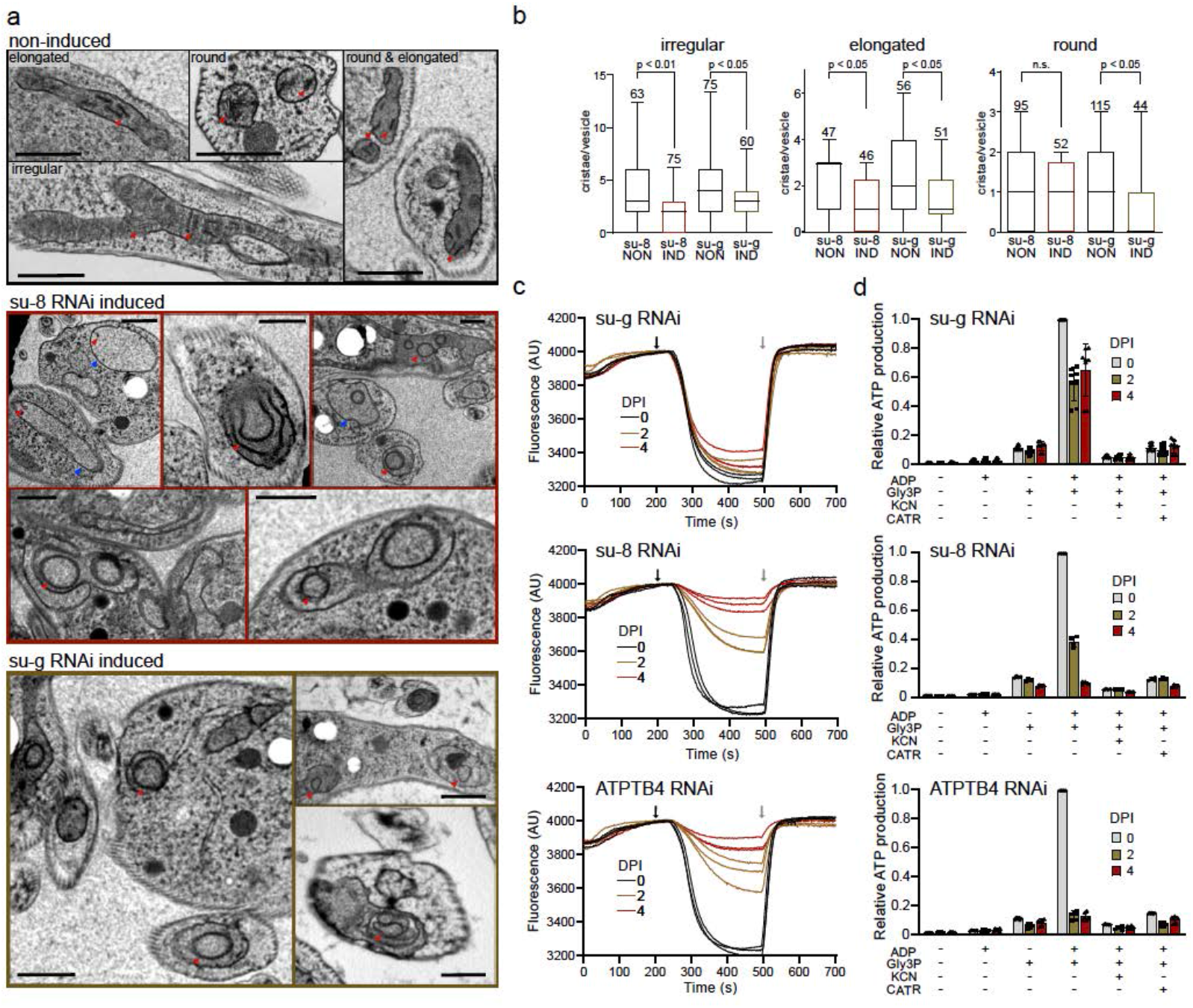
Monomerization of ATP synthase by subunit-*g* knockdown results in aberrant mitochondrial ultrastructure but does not abolish catalytic activity. **a**, Transmission electron micrographs of sections of non-induced or 4 days induced RNAi cell lines. Mitochondrial membranes and cristae are marked with blue and red arrowheads, respectively. Top panel shows examples of irregular, elongated and round cross-sections of mitochondria quantified in (b). **b**, Cristae numbers per vesicle from indicated induced (IND) or non-induced (NON) cell lines counted separately in irregular, elongated and round mitochondrial cross-section. Boxes and whiskers show 25^th^ to 75^th^ and 5^th^ to 95^th^ percentiles, respectively. The numbers of analysed cross-sections are indicated for each data point. Unpaired t-test, p-values are shown. **c**, Mitochondrial membrane polarization capacity of non-induced or RNAi-induced cell lines two and four DPI measured by Safranine O. Black and gray arrow indicate addition of ATP and oligomycin, respectively. **d**, ATP production in permeabilized non-induced (0) or RNAi-induced cells 2 and 4 DPI in the presence of indicated substrates and inhibitors. The graphs show individual values, means (bars) and SD (error bars) of at least four replicates. *Gly3P*, DL-glycerol phosphate; *KCN*, potassium cyanide; *CATR*, carboxyatractyloside

Despite the altered mitochondrial ultrastructure, the subunit-*g*^RNAi^ cells showed only a very mild growth phenotype, in contrast to all other RNAi cell lines that exhibited steadily slowed growth from day three to four after the RNAi induction (Fig. 7a, Extended Data Fig. 9a). This is consistent with the growth defects observed after the ablation of F_o_ subunit ATPTB1^19^ and F_1_ subunits-α and p18^12^. Thus, the monomerization of ATP synthase upon subunit-*g* ablation had only a negligible effect on the fitness of trypanosomes cultured in glucose-rich medium, in which ATP production by substrate level phosphorylation partially compensates for compromised oxidative phosphorylation^37^.

Measurement of oligomycin-sensitive ATP-dependent mitochondrial membrane polarization by safranin O assay in permeabilized cells showed that the proton pumping activity of the ATP synthase in the induced subunit-*g*^RNAi^ cells is negligibly affected, demonstrating that the monomerized enzyme is catalytically functional. By contrast, RNAi downregulation of subunit-8, ATPTB4 and ATPTB11, and ATPTB1 resulted in a strong decline of the mitochondrial membrane polarization capacity, consistent with the loss of both monomeric and dimeric ATP synthase forms (Fig. 7c). Accordingly, knockdown of the same subunits resulted in inability to produce ATP by oxidative phosphorylation (Fig. 7d). However, upon subunit-*g* ablation the ATP production was affected only partially, confirming that the monomerized ATP synthase remains catalytically active. The ~50 % drop in ATP production of subunit-*g*^RNAi^ cells can be attributed to the decreased oxidative phosphorylation efficiency due to the impaired cristae morphology. Indeed, when cells were cultured in the absence of glucose, enforcing the need for oxidative phosphorylation, knockdown of subunit-*g* results in a growth arrest, albeit one to two days later than knockdown of all other tested subunits (Fig. 6a). The data show that dimerization is critical when oxidative phosphorylation is the predominant source of ATP.

## Discussion

Our structure of the mitochondrial ATP synthase dimer from the mammalian parasite *T. brucei* offers new insight into the mechanism of membrane shaping, rotary catalysis, and proton transfer. Considering that trypanosomes belong to an evolutionarily divergent group of Kinetoplastida, the ATP synthase dimer has several interesting features that differ from other dimer structures. The subunit-*b* found in bacterial and other mitochondrial F-type ATP synthases appears to be highly reduced to a single transmembrane helix *b*H1. The long *b*H2, which constitutes the central part of the peripheral stalk in other organisms, and is also involved in the composition of the lumenal proton half-channel, is completely absent in *T. brucei*. Interestingly, the position of *b*H2 in the proton half channel is occupied by a fully ordered phosphatidylcholine molecule that replaces a well-conserved proteinaceous element in the proton path. However, this replacement is not a common trait of all type-IV ATP synthases, because subunit-*b* in *Euglena gracilis* contains the canonical bH2 but lacks bH1^10^. Thus, while subunit-*b* is conserved in Euglenozoa, the lineages of *T. brucei* and *E. gracilis* retained its different non-overlapping structural elements (Fig. 2c). Lack of *b*H2 in *T. brucei* also affects composition of the peripheral stalk in which the divergent subunit-*d* and subunit-*8* binds directly to a C-terminal extension of OSCP, indicating a remodeled peripheral stalk architecture. The peripheral stalk contacts the F_1_ headpiece at several positions conferring greater conformational flexibility to the ATP synthase.

Using the structural and functional data, we also identified a conserved structural element of the ATP synthase that is responsible for its multimerization. Particularly, subunit-*g* is required for the dimerization, but dispensable for the assembly of the F_1_F_o_ monomers. Although the monomerized enzyme is catalytically competent, the inability to form dimers results in defective cristae structure, and consequently leads to compromised oxidative phosphorylation and cease of proliferation. The cristae-shaping properties of mitochondrial ATP synthase dimers are critical for sufficient ATP production by oxidative phosphorylation, but not for other mitochondrial functions, as demonstrated by the lack of growth phenotype of subunit-*g*^RNAi^ cells in the presence of glucose. Thus, trypanosomal subunit-*g* depletion strain represents an experimental tool to assess the roles of the enzyme’s primary catalytic function and mitochondria-specific membrane-shaping activity, highlighting the importance of the latter for oxidative phosphorylation.

Based on our data and previously published structures, we propose an ancestral state with double rows of ATP synthase monomers connected by *e/g* modules longitudinally and by other F_o_ subunits transversally. During the course of evolution, different pairs of adjacent ATP synthase monomer units formed stable dimers in individual lineages (Fig. 8). This gave rise to the highly divergent type-I and type-IV ATP synthase dimers with subunit-*e/g* modules serving either as oligomerization or dimerization motives, respectively. Because trypanosomes belong to the deep-branching eukaryotic supergroup Discoba, the proposed arrangement might have been present in the last eukaryotic common ancestor. Although sequence similarity of subunit-*g* is low and restricted to the single transmembrane helix, we found homologs of subunit-*g* in addition to Opisthokonta and Discoba also in Archaeplastida and Amoebozoa, which represent other eukaryotic supergroups, thus supporting the ancestral role in oligomerization (Extended Data Fig. 8). Taken together, our analysis reveals that mitochondrial ATP synthases that display markedly diverged architecture share the ancestral structural module that promotes oligomerization.

**Fig. 8:**
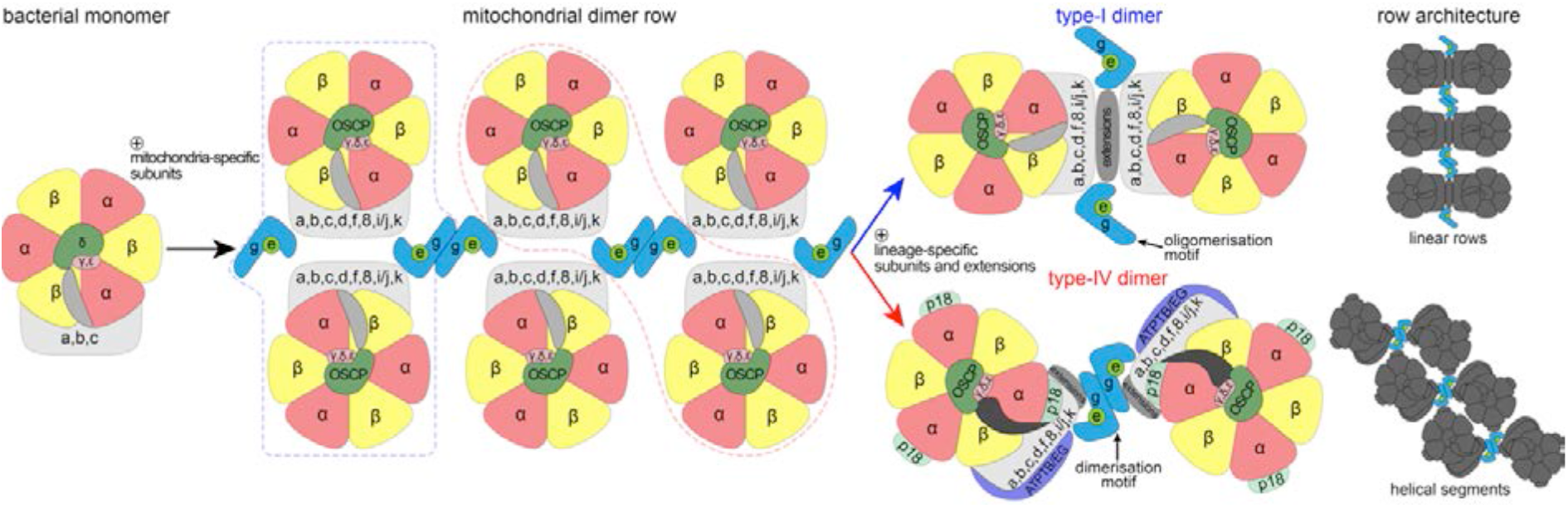
The subunit-*e/g* module is an ancestral oligomerization motif of ATP synthase. Schematic model of the evolution of type-I and IV ATP synthases. Mitochondrial ATP synthases are derived from a monomeric complex of proteobacterial origin. In a mitochondrial ancestor, acquisition of mitochondria-specific subunits, including the subunit-*e/g* module resulted in the assembly of ATP synthase double rows, the structural basis for cristae biogenesis. Through divergence, different ATP synthase architectures evolved, with the subunit-*e/g* module functioning as an oligomerization (type I) or dimerization (type IV) motif, resulting in distinct row assemblies between mitochondrial lineages.

## Materials and Methods

### Cell culture and isolation of mitochondria

*T. brucei* procyclic strains were cultured in SDM-79 medium supplemented with 10% (v/v) fetal bovine serum. For growth curves in glucose-free conditions, cells were grown in SDM-80 medium with 10 % dialysed FBS. RNAi cell lines were grown in presence of 2.5 μg/ml phleomycin and 1 μg/ml puromycin. For ATP synthase purification, mitochondria were isolated from the Lister strain 427. Typically, 1.5×10^11^ cells were harvested, washed in 20 mM sodium phosphate buffer pH 7.9 with 150 mM NaCl and 20 mM glucose, resuspended in hypotonic buffer 1 mM Tris-HCl pH 8.0, 1 mM EDTA, and disrupted by 10 strokes in a 40-ml Dounce homogenizer. The lysis was stopped by immediate addition of sucrose to 0.25 M. Crude mitochondria were pelleted (15 min at 16,000 xg, 4°C), resuspended in 20 mM Tris-HCl pH 8.0, 250 mM sucrose, 5 mM MgCl_2_, 0.3 mM CaCl_2_ and treated with 5 μg/ml DNase I. After 60 min on ice, one volume of the STE buffer (20 mM Tris-HCl pH 8.0, 250 mM sucrose, 2 mM EDTA) was added and mitochondria were pelleted (15 min at 16000 xg, 4°C). The pellet was resuspended in 60% (v/v) Percoll in STE and loaded on six linear 10-35% Percoll gradients in STE in polycarbonate tubes for SW28 rotor (Beckman). Gradients were centrifuged for 1 h at 24,000 rpm, 4°C. The middle phase containing mitochondrial vesicles (15-20 ml per tube) was collected, washed four times in the STE buffer, and pellets were snap-frozen in liquid nitrogen and stored at −80°C.

### Plasmid construction and generation of RNAi cell lines

To downregulate ATP synthase subunits by RNAi, DNA fragments corresponding to individual target sequences were amplified by PCR from Lister 427 strain genomic DNA using forward and reverse primers extended with restriction sites *Xho*I&*Kpn*I and *Xba*I&*BamH*I, respectively (Extended Data Table 3). Each fragment was inserted into the multiple cloning sites 1 and 2 of pAZ0055 vector, derived from pRP^HYG-iSL^ (courtesy of Sam Alsford) by replacement of hygromycine resistance gene with phleomycine resistance gene, with restriction enzymes *Kpn*I/*Bam*HI and *Xho*I/*Xba*I, respectively. Resulting constructs with tetracycline inducible T7 polymerase driven RNAi cassettes were linearized with *NotI* and transfected into a cell line derived from the Lister strain 427 by integration of the SmOx construct for expression of T7 polymerase and the tetracycline repressor^38^ into the β-tubulin locus. RNAi was induced in selected semi-clonal populations by addition of 1 μg/ml tetracycline and the downregulation of target mRNAs was verified by quantitative RT-PCR 2 and 4 days post induction. The total RNA isolated by an RNeasy Mini Kit (Qiagen) was treated with 2 μg of DNase I, and then reverse transcribed to cDNA with TaqMan Reverse Transcription kit (Applied Biosciences). qPCR reactions were set with Light Cycler 480 SYBR Green I Master mix (Roche), 2 μl of cDNA and 0.3 μM primers (Extended Data Table 3), and run on LightCycler 480 (Roche). Relative expression of target genes was calculated using - ΔΔCt method with 18S rRNA or β-tubulin as endogenous reference genes and normalized to noninduced cells.

### Denaturing and blue native polyacrylamide electrophoresis and immunoblotting

Whole cell lysates for denaturing sodium dodecyl sulphate polyacrylamide electrophoresis (SDS-PAGE) were prepared from cells resuspended in PBS buffer (10 mM phosphate buffer, 130 mM NaCl, pH 7.3) by addition of 3x Laemmli buffer (150 mM Tris pH 6.8, 300 mM 1,4-dithiothreitol, 6% (w/v) SDS, 30% (w/v) glycerol, 0.02% (w/v) bromophenol blue) to final concentration of 1×10^7^ cells in 30 μl. The lysates were boiled at 97°C for 10 min and stored at −20°C. For immunoblotting, lysates from 3×10^6^ cells were separated on 4-20 % gradient Tris-glycine polyacrylamide gels (BioRad 4568094), electroblotted onto a PVDF membrane (Pierce 88518), and probed with respective antibodies (Extended Data Table 4). Membranes were incubated with the Clarity Western ECL substrate (BioRad 1705060EM) and chemiluminescence was detected on a ChemiDoc instrument (BioRad). Band intensities were quantified densitometrically using the ImageLab software. The levels of individual subunits were normalized to the signal of mtHsp70.

Blue native PAGE (BN-PAGE) was performed as described earlier^12^ with following modifications. Crude mitochondrial vesicles from 2.5 × 10^8^ cells were resuspended in 40 μl of Solubilization buffer A (2 mM ε-aminocaproic acid (ACA), 1 mM EDTA, 50 mM NaCl, 50 mM Bis-Tris/HCl, pH 7.0) and solubilized with 2% (w/v) dodecylmaltoside (β-DDM) for 1 h on ice. Lysates were cleared at 16,000 g for 30 min at 4°C and their protein concentration was estimated using bicinchoninic acid assay. For each time point, a volume of mitochondrial lysate corresponding to 4 μg of total protein was mixed with 1.5 μl of loading dye (500 mM ACA, 5% (w/v) Coomassie Brilliant Blue G-250) and 5% (w/v) glycerol and with 1 M ACA until a final volume of 20 μl/well, and resolved on a native PAGE 3-12% Bis-Tris gel (Invitrogen). After the electrophoresis (3 h, 140 V, 4°C), proteins were transferred by electroblotting onto a PVDF membrane (2 h, 100 V, 4°C, stirring), followed by immunodetection with an appropriate antibody (Extended Data Table 4).

### Mitochondrial membrane polarization measurement

The capacity to polarize mitochondrial membrane was determined fluorometrically employing safranin O dye (Sigma S2255) in permeabilized cells. For each sample, 2×10^7^ cells were harvested and washed with ANT buffer (8 mM KCl, 110 mM K-gluconate, 10 mM NaCl, 10 mM free-acid Hepes, 10 mM K_2_HPO_4_, 0.015 mM EGTA potassium salt, 10 mM mannitol, 0.5 mg/ml fatty acid-free BSA, 1.5 mM MgCl_2_, pH 7.25). The cells were permeabilized by 8 μM digitonin in 2 ml of ANT buffer containing 5 μM safranin O. Fluorescence was recorded for 700 s in a Hitachi F-7100 spectrofluorimeter (Hitachi High Technologies) at a 5-Hz acquisition rate, using 495 nm and 585 nm excitation and emission wavelengths, respectively. 1 mM ATP (PanReac AppliChem A1348,0025) and 10 μg/ml oligomycin (Sigma O4876) were added after 230 s and 500 s, respectively. Final addition of the uncoupler SF 6847 (250 nM; Enzo Life Sciences BML-EI215-0050) served as a control for maximal depolarization. All experiments were performed at room temperature and constant stirring.

### ATP production assay

ATP production in digitonin-isolated mitochondria was performed as described previously^39^. Briefly, 1×10^8^ cells per time point were lysed in SoTE buffer (600 mM sorbitol, 2 mM EDTA, 20 mM Tris-HCl, pH 7.75) containing 0.015% (w/v) digitonin for 5 min on ice. After centrifugation (3 min, 4,000 g, 4°C), the soluble cytosolic fraction was discarded and the organellar pellet was resuspended in 75 μl of ATP production assay buffer (600 mM sorbitol, 10 mM MgSO_4_, 15 mM potassium phosphate buffer pH 7.4, 20 mM Tris-HCl pH 7.4, 2.5 mg/ml fatty acid-free BSA). ATP production was induced by addition of 20 mM DL-glycerol phosphate (sodium salt) and 67 μM ADP. Control samples were preincubated with the inhibitors potassium cyanide (1 mM) and carboxyatractyloside (6.5 μM) for 10 min at room temperature. After 30 min at room temperature, the reaction was stopped by addition of 1.5 μl of 70% perchloric acid. The concentration of ATP was estimated using the Roche ATP Bioluminescence Assay Kit HS II in a Tecan Spark plate reader. The luminescence values of the RNAi induced samples were normalized to that of the corresponding noninduced sample.

### Thin sectioning and transmission electron microscopy

The samples were centrifuged and pellet was transferred to the specimen carriers which were completed with 20% BSA and immediately frozen using high pressure freezer Leica EM ICE (Leica Microsystems). Freeze substitution was performed in the presence of 2% osmium tetroxide diluted in 100% acetone at −90°C. After 96 h, specimens were warmed to −20°C at a slope 5 °C/h. After the next 24 h, the temperature was increased to 3°C (3°C/h). At room temperature, samples were washed in acetone and infiltrated with 25%, 50%, 75% acetone/resin EMbed 812 (EMS) mixture 1 h at each step. Finally, samples were infiltrated in 100% resin and polymerized at 60°C for 48h. Ultrathin sections (70 nm) were cut using a diamond knife, placed on copper grids and stained with uranyl acetate and lead citrate. TEM micrographs were taken with Mega View III camera (SIS) using a JEOL 1010 TEM operating at an accelerating voltage of 80 kV.

### Purification of *T. brucei* ATP synthase dimers

Mitochondria from 3×10^11^ cells were lysed by 1 % (w/v) β-DDM in 60 ml of 20 mM Bis-tris propane pH 8.0 with 10 % glycerol and EDTA-free Complete protease inhibitors (Roche) for 20 min at 4°C. The lysate was cleared by centrifugation at 30,000 xg for 20 min at 4°C and adjusted to pH 6.8 by drop-wise addition of 1 M 3-(N-morpholino) propanesulfonic acid pH 5.9. Recombinant TbIF_1_ without dimerization region, whose affinity to F_1_-ATPase was increased by N-terminal truncation and substitution of tyrosine 36 with tryptophan^20^, with a C-terminal glutathione S-transferase (GST) tag (TbIF_1_(9-64)-Y36W-GST) was added in approximately 10-fold molar excess over the estimated content of ATP synthase. Binding of TbIF_1_ was facilitated by addition of neutralized 2 mM ATP with 4 mM magnesium sulphate. After 5 min, sodium chloride was added to 100 mM, the lysate was filtered through a 0.2 μm syringe filter and immediately loaded on 5 ml GSTrap HP column (Cytiva) equilibrated in 20 mM Bis-Tris-Propane pH 6.8 binding buffer containing 0.1 % (w/v) glyco-diosgenin (GDN; Avanti Polar Lipids), 10 % (v/v) glycerol, 100 mM sodium chloride, 1 mM tris(2-carboxyethyl)phosphine (TCEP), 1 mM ATP, 2 mM magnesium sulphate, 15 μg/ml cardiolipin, 50 μg/ml 1-palmitoyl-2-oleoyl-sn-glycero-3-phosphocholine (POPC), 25 μg/ml 1-palmitoyl-2-oleoyl-sn-glycero-3-phosphoethanolamine (POPE) and 10 μg/ml 1-palmitoyl-2-oleoyl-sn-glycero-3-[phospho-rac-(1-glycerol)] (POPG). All phospholipids were purchased from Avanti Polar Lipids (catalog numbers 840012C, 850457C, 850757C and 840757, respectively). ATP synthase was eluted with a gradient of 20 mM reduced glutathione in Tris pH 8.0 buffer containing the same components as the binding buffer. Fractions containing ATP synthase were pooled and concentrated to 150 μl on Vivaspin centrifugal concentrator with 30 kDa molecular weight cut-off. The sample was fractionated by size exclusion chromatography on a Superose 6 Increase 3.2/300 GL column (Cytiva) equilibrated in a buffer containing 20 mM Tris pH 8.0, 100 mM sodium chloride, 2 mM magnesium chloride, 0.1 % (w/v) GDN, 3.75 μg/ml cardiolipin, 12.5 μg/ml POPC, 6.25 μg/ml POPE and 2.5 μg/ml POPG at 0.03 ml/min. Fractions corresponding to ATP synthase were pooled, supplemented with 0.05% (w/v) β-DDM that we and others experimentally found to better preserve dimer assemblies in cryo-EM^40^, and concentrated to 50 μl.

### Preparation of cryo-EM grids and data collection

Samples were vitrified on glow-discharged Quantifoil R1.2/1.3 Au 300-mesh grids after blotting for 3 sec, followed by plunging into liquid ethane using a Vitrobot Mark IV. 5,199 movies were collected using EPU 1.9 on a Titan Krios (ThermoFisher Scientific) operated at 300kV at a nominal magnification of 165 kx (0.83 Å/pixel) with a Quantum K2 camera (Gatan) using a slit width of 20 eV. Data was collected with an exposure rate of 3.6 electrons/px/s, a total exposure of 33 electrons/Å^2^ and 20 frames per movie.

### Image processing

Image processing was performed within the Scipion 2 framework^41^, using RELION-3.0 unless specified otherwise. Movies were motion-corrected using the RELION implementation of the MotionCor2. 294,054 particles were initially picked using reference-based picking in Gautomatch (http://www.mrc-lmb.cam.ac.uk/kzhang/Gautomatch) and Contrast-transfer function parameters were using GCTF^42^. Subsequent image processing was performed in RELION-3.0 and 2D and 3D classification was used to select 100,605 particles, which were then extracted in an unbinned 560-pixel box (Fig. S1). An initial model of the ATP synthase dimer was obtained using *de novo* 3D model generation. Using masked refinement with applied C2 symmetry, a 2.7-Å structure of the membrane region was obtained following per-particle CTF refinement and Bayesian polishing. Following C2-symmetry expansion and signal subtraction of one monomer, a 3.7 Å map of the peripheral stalk was obtained. Using 3D classification (T=100) of aligned particles, with a mask on the F_1_/*c*-ring region, 10 different rotational substates were then separated and maps at 3.5-4.3 Å resolution were obtained using 3D refinement. The authors note that the number of classes identified in this study likely reflects the limited number of particles, rather than the complete conformational space of the complex. By combining particles from all states belonging to main rotational state 1, a 3.7-Å map of the rotor and a 3.2-Å consensus map of the complete ATP synthase dimer with both rotors in main rotational state 1 were obtained.

### Model building, refinement and data visualization

An initial atomic model of the static F_o_ membrane region was built automatically using Bucaneer^43^. Subunits were subsequently assigned directly from the cryo-EM map, 15 of them corresponding to previously identified *T. brucei* ATP synthase subunits^21^, while three subunits (ATPTB14, ATPEG3, ATPEG4) were newly identified using BLAST searches. Manual model building was performed in *Coot* using the *T. brucei* F_1_ (PDB 6F5D)^13^ and homology models^44^ of the *E. gracilis* OSCP and *c*-ring (PDB 6TDU)^10^ as starting models. Ligands were manually fitted to the map and restraints were generated by the GRADE server (http://grade.globalphasing.org). Cardiolipins were assigned based on the presence of a characteristic elongated density branched on both termini, corresponding to two phosphatidyl groups linked by the central glycerol bridge. Monophosphatidyl lipids were assigned based on their headgroup densities. Characteristic tetrahedral shapes of densities of choline groups served to distinguish phosphatidylcholines from elongated phosphatidylethanolamine head groups (Extended Data Figure 5g,h). Real-space refinement was performed in PHENIX using auto-sharpened, local-resolution-filtered maps of the membrane region, peripheral stalk tip, *c*-ring/central stalk and F_1_F_o_ monomers in different rotational states, respectively, using secondary structure restraints. Model statistics were generated using MolProbity^45^ and EMRinger^46^ Finally, the respective refined models were combined into a composite ATP synthase dimer model and real-space refined against the local-resolution-filtered consensus ATP synthase dimer map with both monomers in rotational state 1, applying reference restraints. Figures of the structures were prepared using ChimeraX^47^, the proton half-channels were traced using HOLLOW^48^.

## Data availability

The atomic coordinates have been deposited in the Protein Data Bank (PDB) and are available under the accession codes: XXXX (membrane-region), XXXX (peripheral stalk), XXXX (rotor), XXXX (F_1_F_o_ dimer), XXXX (rotational state 1a), XXXX (rotational state 1b), XXXX (rotational state 1c), XXXX (rotational state 1d), XXXX (rotational state 1e), XXXX (rotational state 2a), XXXX (rotational state 2b), XXXX (rotational state 2c), XXXX (rotational state 2d), XXXX (rotational state 3). The local resolution filtered cryo-EM maps, half maps, masks and FSC-curves have been deposited in the Electron Microscopy Data Bank with the accession codes: EMD-XXXX (membrane-region), EMD-XXXX (peripheral stalk), EMD-XXXX (rotor), EMD-XXXX (F_1_F_o_ dimer), EMD-XXXX (rotational state 1a), EMD-XXXX (rotational state 1b), EMD-XXXX (rotational state 1c), EMD-XXXX (rotational state 1d), EMD-XXXX (rotational state 1e), EMD-XXXX (rotational state 2a), EMD-XXXX (rotational state 2b), EMD-XXXX (rotational state 2c), EMD-XXXX (rotational state 2d), EMD-XXXX (rotational state 3). Source data are provided with this paper.

## Acknowledgements

We are grateful to Sir John E. Walker and Martin G. Montgomery for invaluable assistance with ATP synthase purification in the initial stage of the project. We acknowledge cryo-electron microscopy and tomography core facility of CIISB, Instruct-CZ Centre, supported by MEYS CR (LM2018127). This work was supported by the Czech Science Foundation grants number 18-17529S to A.Z. and 20-04150Y to O.G. and by European Regional Development Fund (ERDF) and Ministry of Education, Youth and Sport (MEYS) project CZ.02.1.01/0.0/0.0/16_019/0000759 to A.Z., Swedish Foundation for Strategic Research (FFL15:0325), Ragnar Söderberg Foundation (M44/16), European Research Council (ERC-2018-StG-805230), Knut and Alice Wallenberg Foundation (2018.0080), and EMBO Young Investigator Programme to A.A.

## Author contributions

A.Z. and A.A. conceived and designed the work. O.G. prepared the sample for cryo-EM. O.G. and A.M. performed initial screening. A.M. processed the cryo-EM data and built the model. O.G., A.M. and A.A. analyzed the structure. B.P., C.H.Y., M.J., M.S., O.G. and A.Z. performed biochemical analysis. O.G., A.M., A.A. and A.Z. interpreted the data. O.G., A.M., A.A. and A.Z. wrote and revised the manuscript. All authors contributed to the analysis and approved the final version of the manuscript.

---

**Extended Data Fig. 1.**
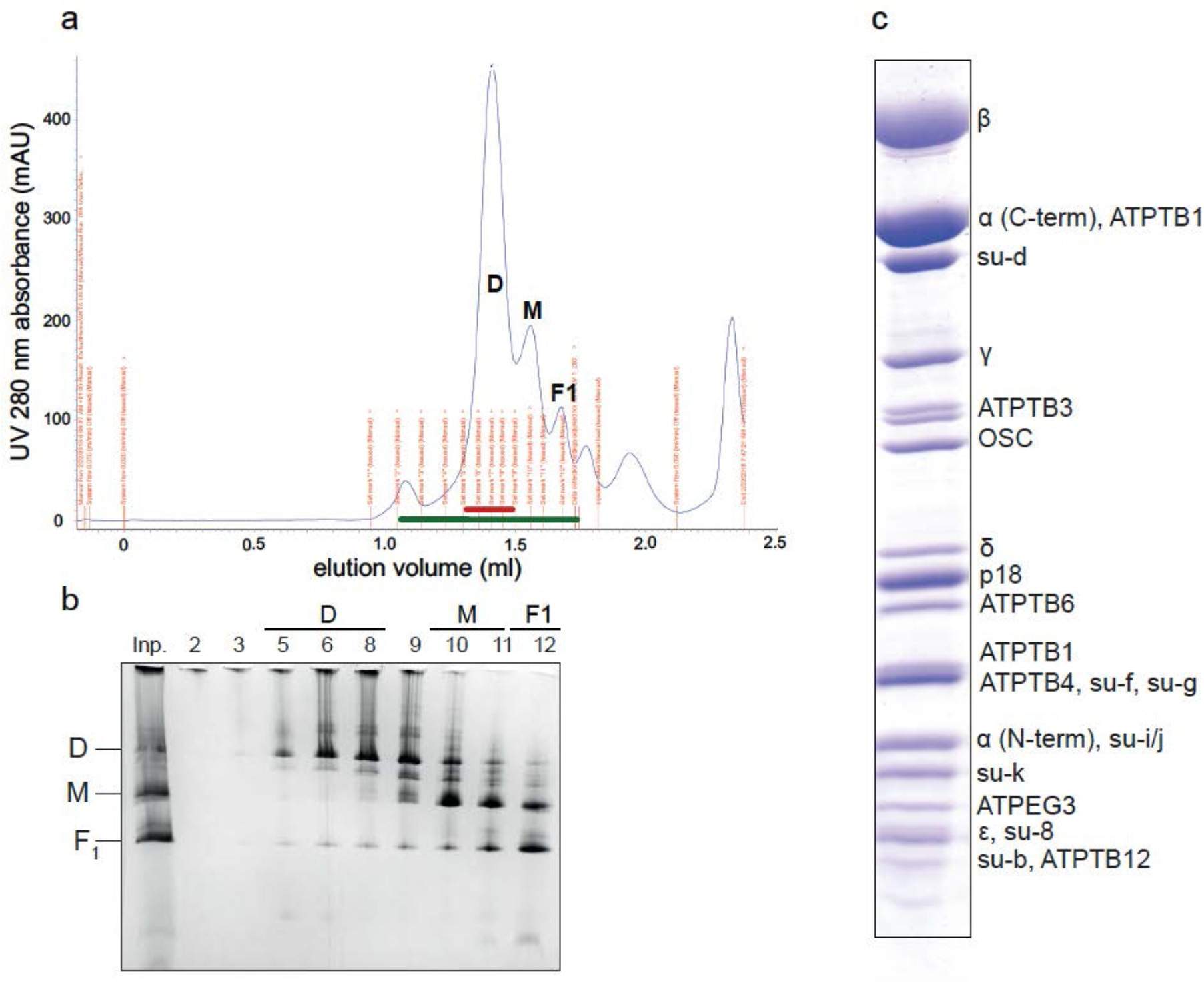
Purification of the *T. brucei* ATP synthase dimer. **a**, Size exclusion chromatography trace with peaks enriched with ATP synthase dimers (D), monomers (M) and F_1_-ATPase (F_1_) labelled. The red bar marks the fractions for cryo-EM. **b,** Fractions from size exclusion chromatography marked with green bar in (a) resolved by native BN-PAGE. **c,** Dimer-enriched fractions resolved by SDS-PAGE stained by Coomassie blue dye. Bands are annotated based on mass spectrometry identification from excised gel pieces.

**Extended Data Fig. 2.**
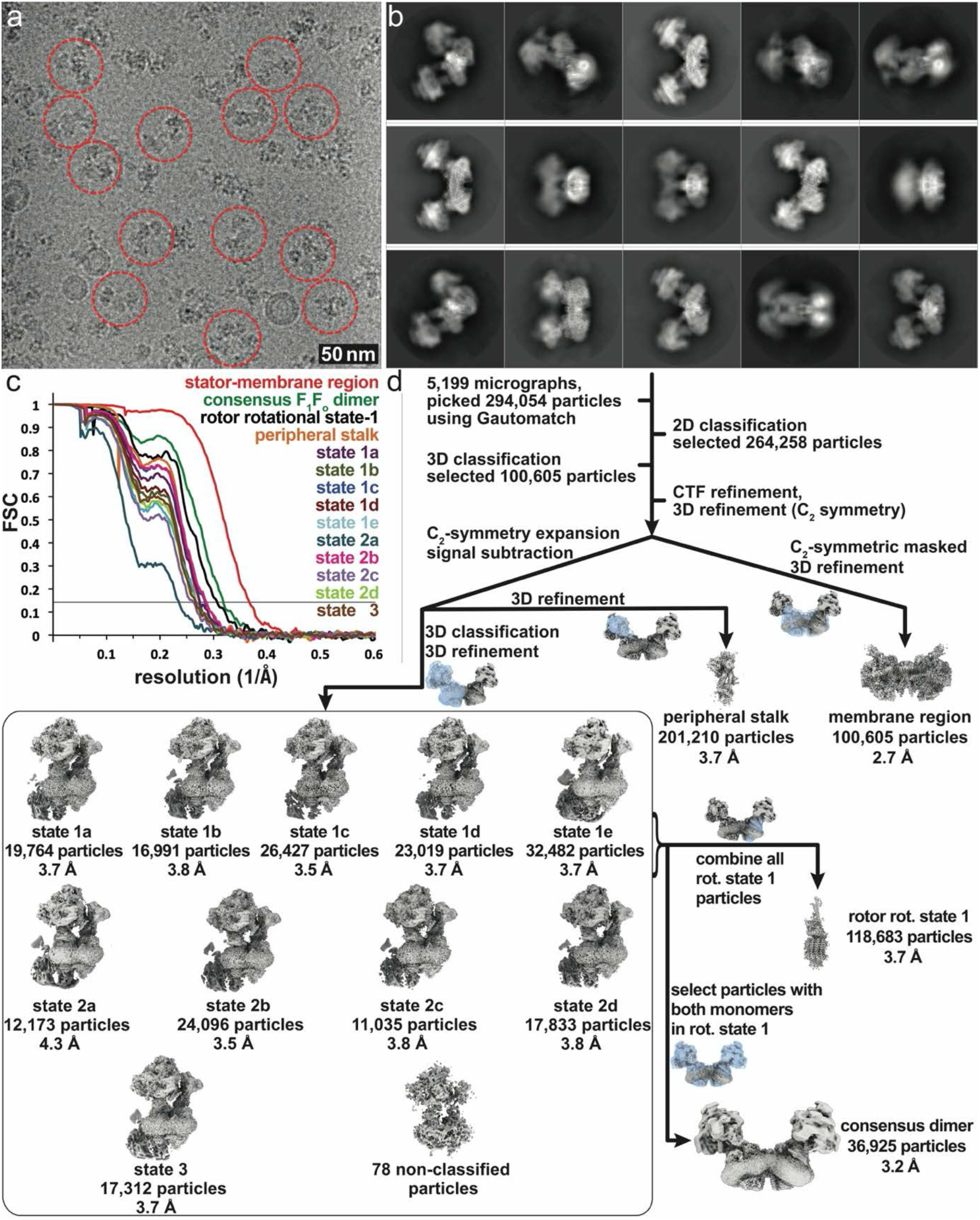
Cryo-EM data processing of the *T. brucei* ATP synthase dimer. **a**, Representative micrograph. **b**, 2D class averages. **c**, Fourier Shell Correlation (FSC) curves showing the estimated resolutions of ATP synthase maps according to the gold standard 0.143 criterion. **d**, Data processing scheme resulting in maps covering all regions of the complex, as well as 10 rotational states.

**Extended Data Fig. 3.**
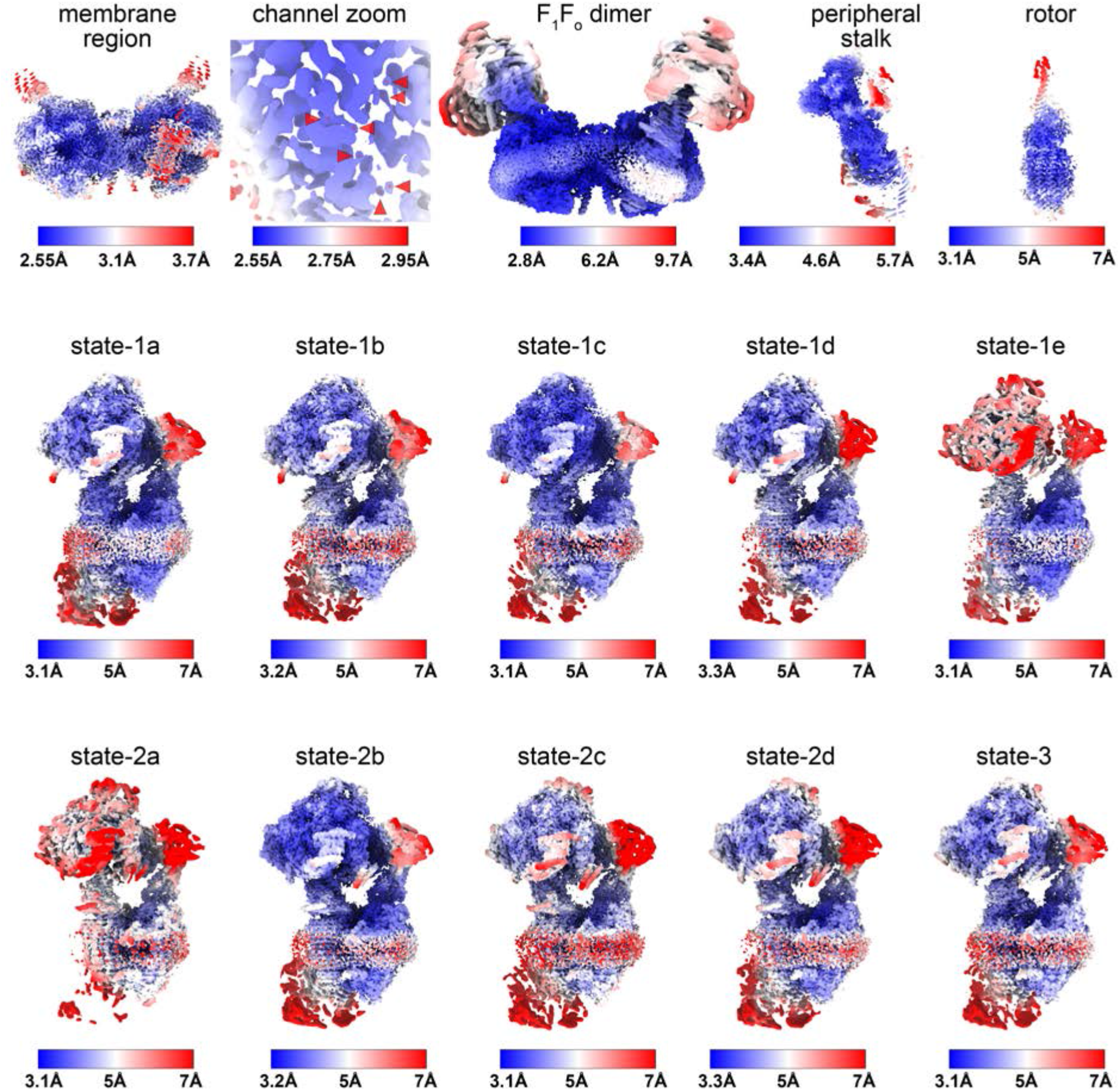
Local resolution estimation of final cryo-em maps. Local resolution estimates colored according to the respective color legends of the membrane region, F_1_F_o_ dimer, the peripheral stalk, the rotor and all identified rotational states. A zoomed-in view of the membrane region shows that the resolution in the lumenal channel extends to 2.55 Å, allowing the assignment of water molecules.

**Extended Data Fig. 4.**
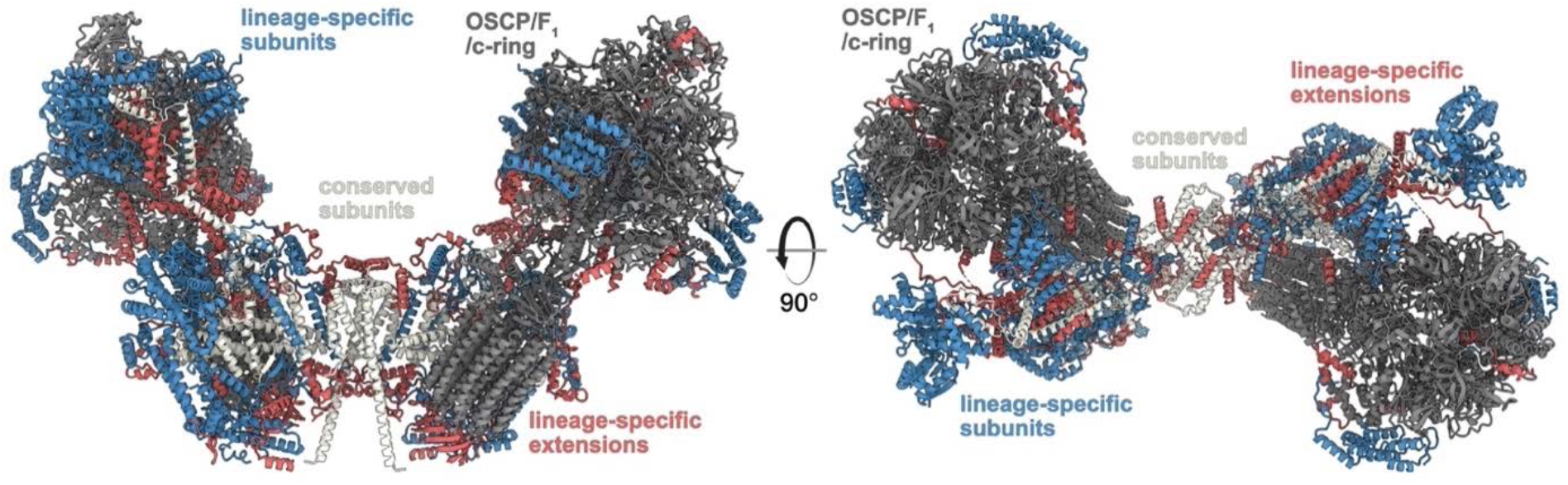
Conserved and phylum specific elements generate the *T. brucei* ATP synthase architecture. The canonical OSCP/F_1_/*c*-ring monomers (dark grey) are tied together by both conserved F_o_ subunits and extensions of lineage-specific subunits (red). The F_o_ periphery and peripheral stalk attachment are composed of lineage specific subunits (blue).

**Extended Data Fig. 5.**
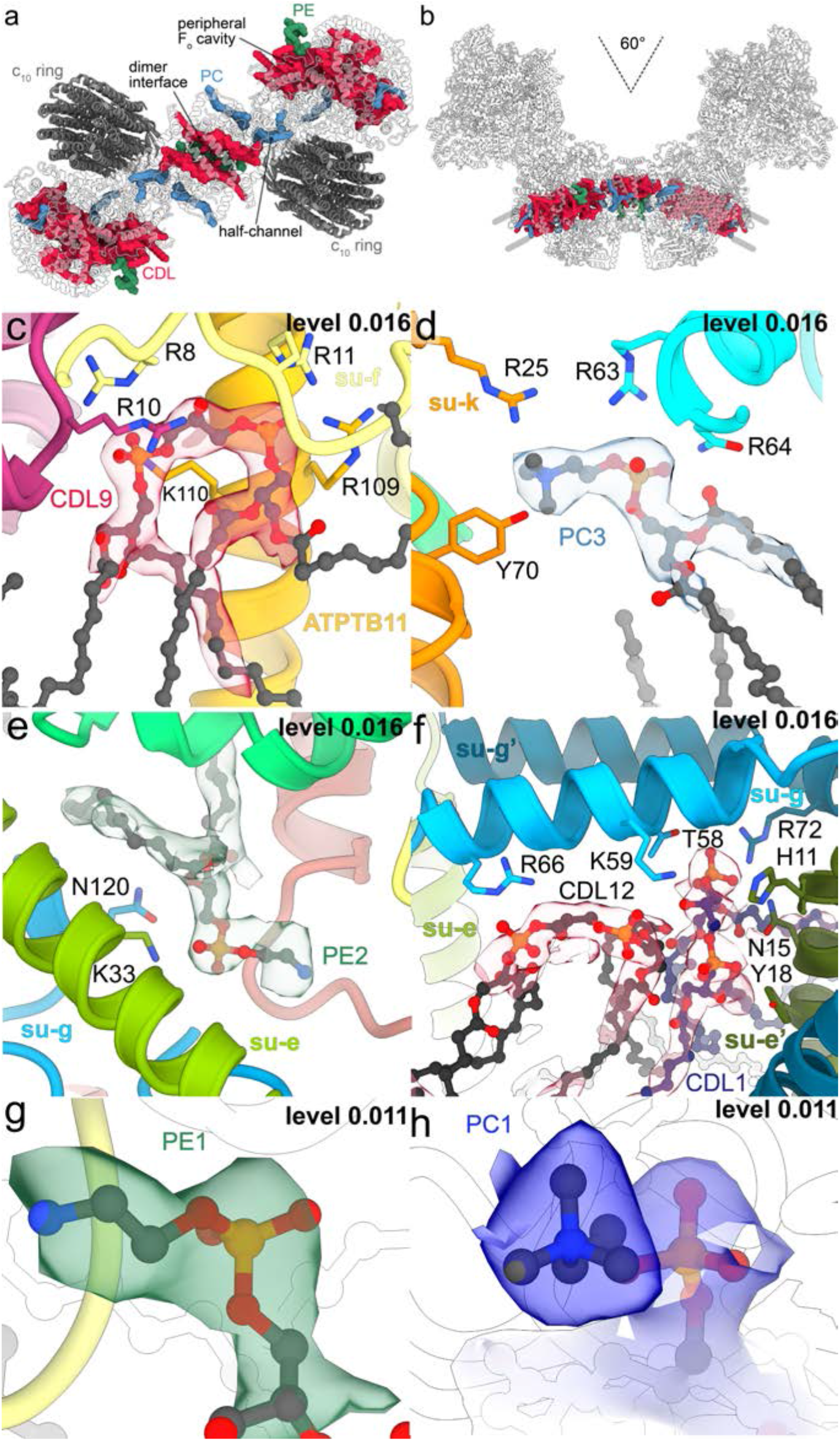
The F_o_ region coordinates numerous bound lipids. **a**, F_o_ top view, cardiolipin (CDL), phosphatidylcholine (PC) and phosphatidlyethanolamine (PE) are bound at the dimer interface, the lumenal proton half-channel and the peripheral F_o_ cavity. **b**, The 60°-dimer angle generates a curved F_o_ region with phospholipids bound in an arc-shaped bilayer. **c-f**, Bound lipids with cryo-EM density and coordinating residues. **g-h,** Representative densities of headgroups of PE (g) and PC (h).

**Extended Data Fig. 6.**
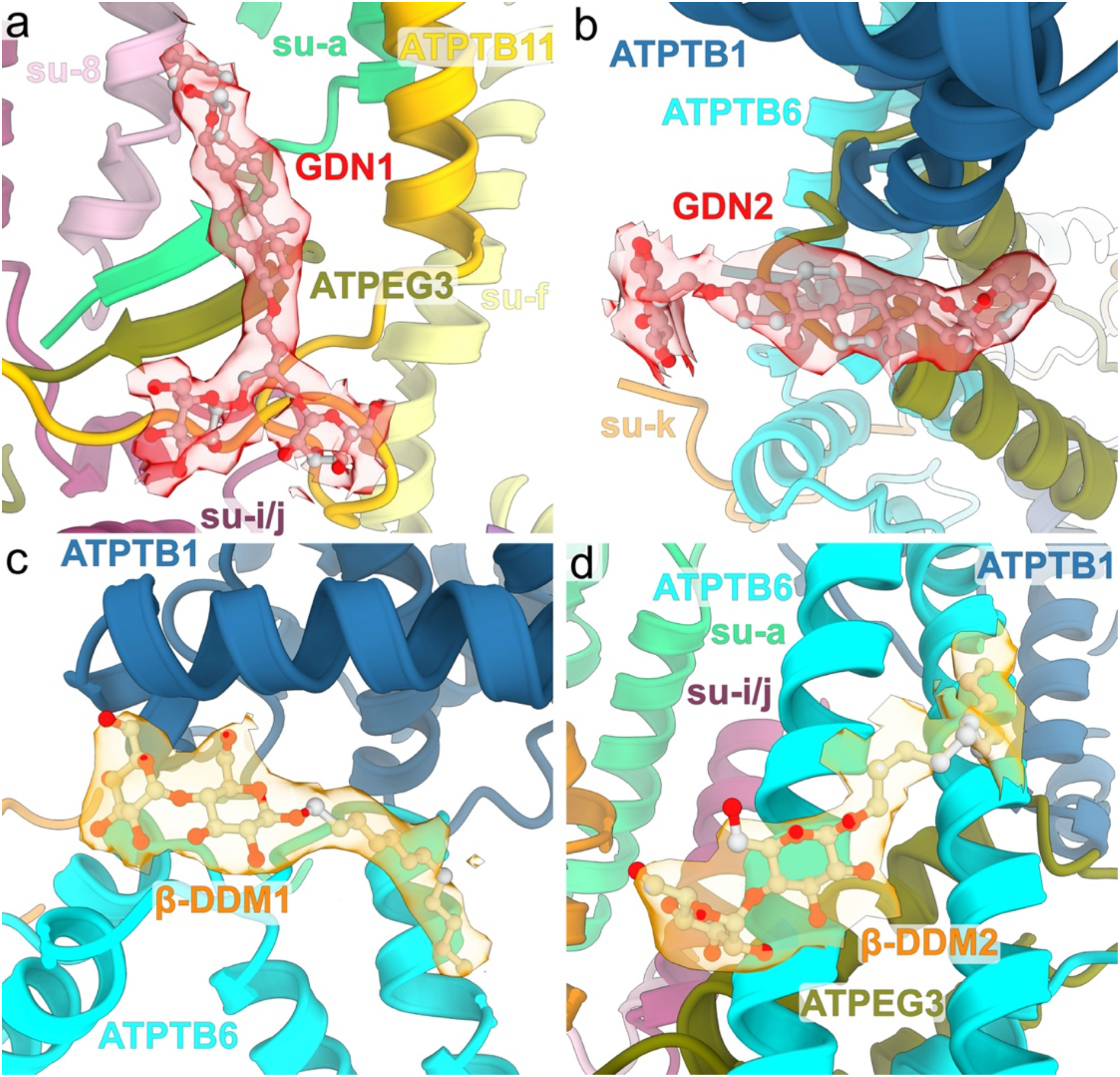
Bound detergents of the F_o_ region. GDN **(a,b)** and β-DDM **(c,d)** molecules bound in the periphery of the membrane region with cryo-EM map densities shown (transparent), indicating that both glycosides are retained in the detergent micelle.

**Extended Data Fig. 7.**
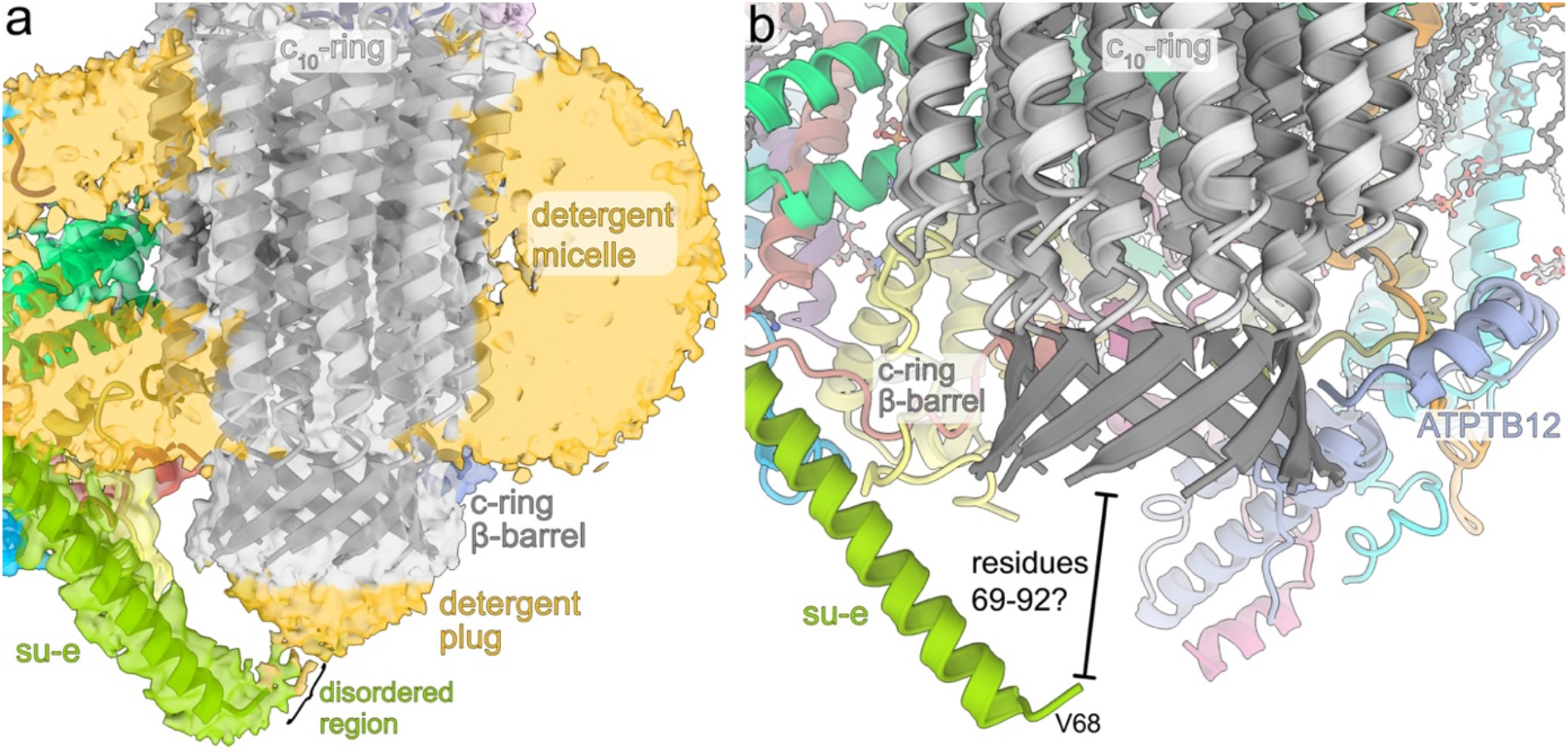
The C-terminal tail of subunit-*e* interacts with the *c*_10_-ring. **a,** The cryo-EM map reveals disordered detergent density of the detergent belt surrounding the membrane region as well as a detergent plug on the luminal side of the *c*-ring. **b,** The helical C-terminus of subunit-*e* extends into the lumen towards the *c*-ring. The terminal 23 residues are disordered and likely interact with the β-barrel.

**Extended Data Fig. 8.**
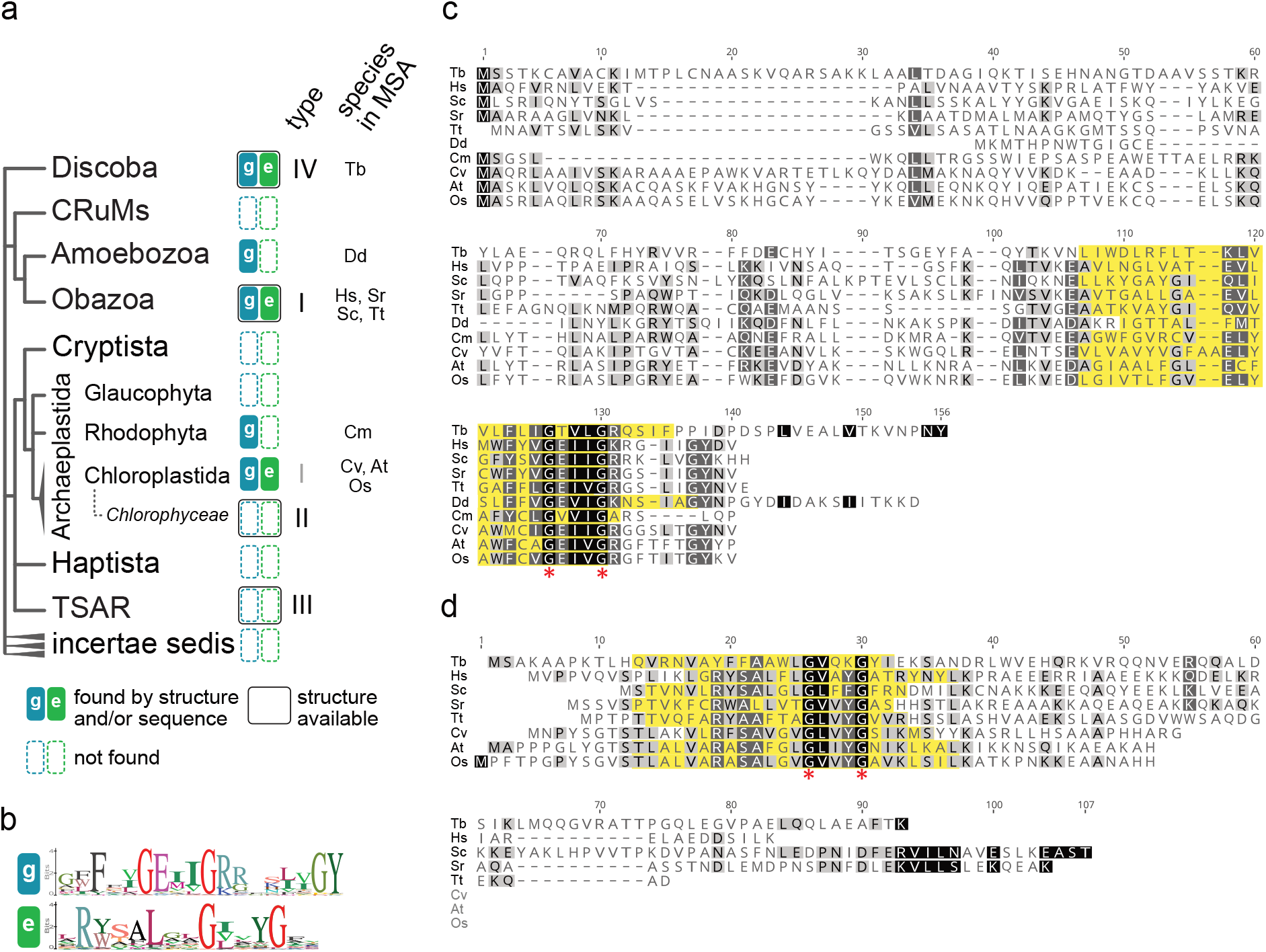
Phylogenetic distribution and sequence conservancy of subunit-*e* and *-g*. **a**, Distribution of subunits *e* and *g* mapped on the phylogenetic tree of eukaryotes^3^. Homologs of subunits *e* and *g* were searched in non-redundant GenBank and UniprotKB protein databases by PSI-BLAST, and phmmer and hmmsearch^4^, respectively, using individual sequences of representatives from *H. sapiens* and *T. brucei*, and in the case of hmmsearch a multiple sequence alignment (MSA) of representatives from *Homo sapiens*, *Saccharomyces cerevisiae*, *Arabidopsis thaliana* and *T. brucei*, as queries. Groups, in which at least one structure of ATP synthase is available, are marked. Abbreviations of species used in MSA in panels (c) and (d) are shown. **b**, Sequence logo of GXXXG motifs and flanking regions of subunits *e* and *g*. Hits from hmmsearch were clustered by CD-HIT Suite^5^ to 50% sequence identity and MSA of representative sequences of each cluster was generated by Clustal Omega4^6^. The sequence logos were created from MSA in Geneious Prime (Biomatters Ltd.). **c,d**, MSA of sequences of subunits g (c) and e (d) from species representing major groups shown in (a) generated by MUSCLE^7^ and visualized in Geneious Prime. The experimentally determined or predicted transmembrane regions are highlighted in yellow. Species abbreviations: Tb – *T. brucei*, Hs – *H. sapiens*, Sc – *S. cerevisiae*, Sr – *Salpingoeca rosetta*, Tt – *Thecamonas trahens*, Dd – *Dictyostelium discoideum*, Cm – *Cyanidioschyzon merolae*, Cv – *Chlorella vulgaris*, At – *Arabidopsis thaliana*, Os – *Oryza sativa*.

**Extended Data Fig. 9.**
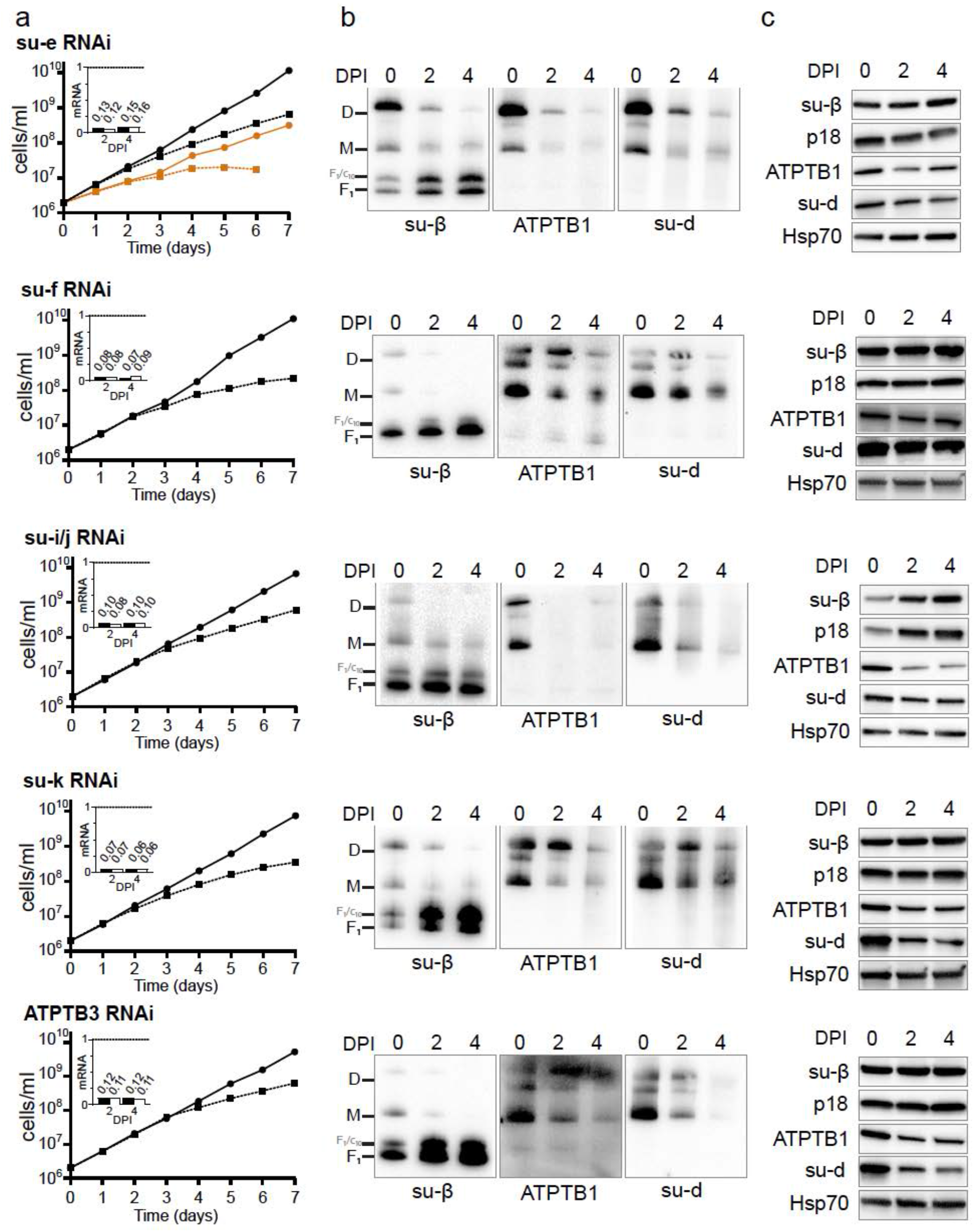
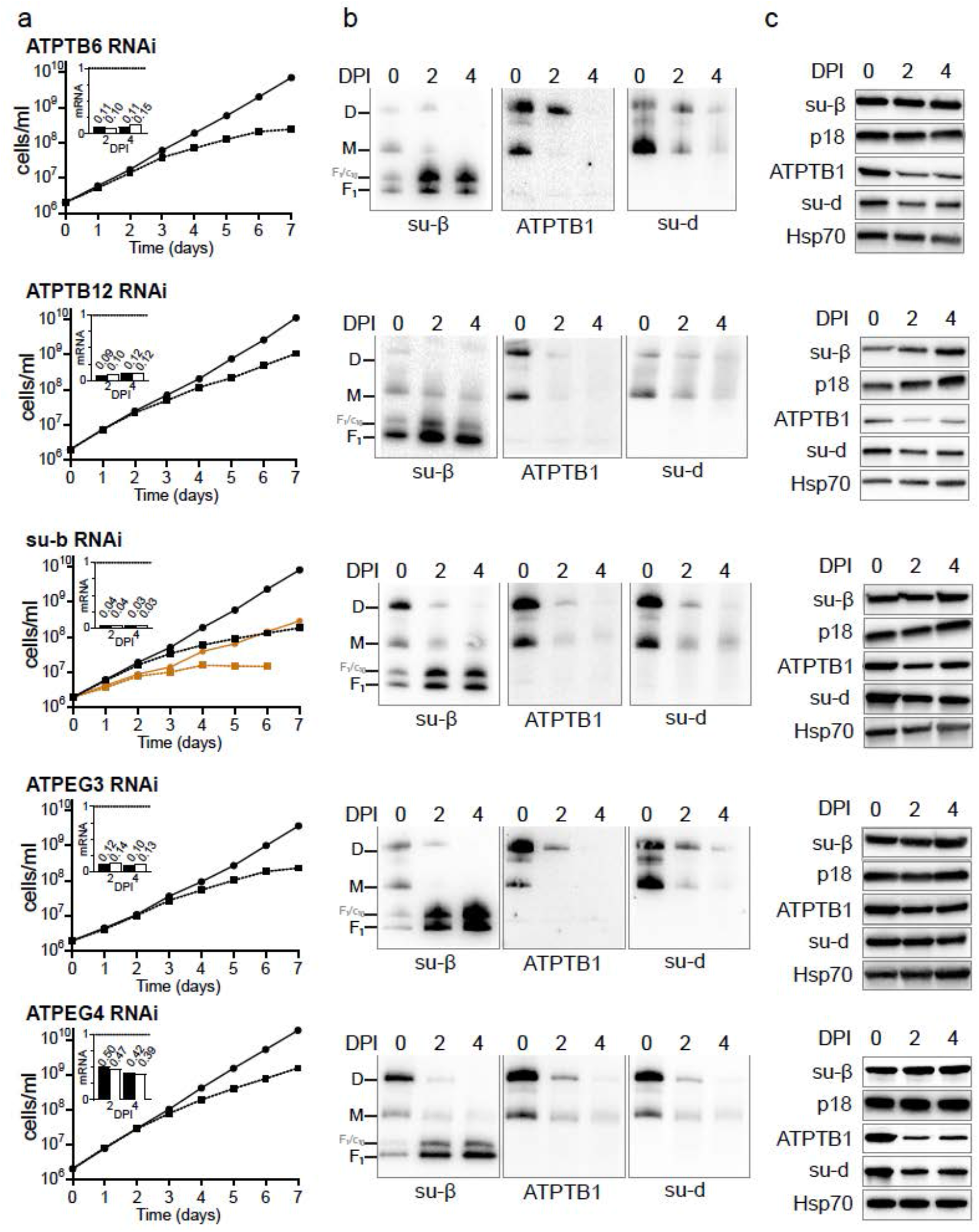
Effects of RNAi knock-down of ATP synthase subunits on viability and stability and dimerization of ATP synthase. **a**, Growth curves of indicated non-induced (solid lines) and tetracycline induced (dashed lines) RNAi cells lines in the presence (black) or absence (brown) of glucose. The insets show relative levels of the respective target mRNA at indicated days post induction (DPI) normalized to the levels of 18S rRNA (black bars) or β-tubulin (white bars). **b**, Immunoblots of mitochondrial lysates from indicated RNAi cell lines resolved by BN-PAGE probed by antibodies against indicated ATP synthase subunits. **c**, Immunoblots of whole cell lysates from indicated RNAi cell lines probed with indicated antibodies.

**Extended Data Table 1.** Data collection, processing, model refinement and validation statistics.

|  | Mem-<br>brane<br>region | Rotor | Periphe-<br>ral stalk | F<sub>1</sub>F<sub>0</sub><br>dimer | Rot.<br>1a | Rot.<br>1b | Rot.<br>1c | Rot.<br>1d | Rot.<br>1e | Rot.<br>2a | Rot.<br>2b | Rot.<br>2c | Rot.<br>2d | Rot.<br>3 |
| --- | --- | --- | --- | --- | --- | --- | --- | --- | --- | --- | --- | --- | --- | --- |
| Data collection |  |  |  |  |  |  |  |  |  |  |  |  |  |  |
| Microscope | Titan Krios |  |  |  |  |  |  |  |  |  |  |  |  |  |
| Voltage (kV) | 300 |  |  |  |  |  |  |  |  |  |  |  |  |  |
| Camera | K2 Summit |  |  |  |  |  |  |  |  |  |  |  |  |  |
| Magnification | 165 kx |  |  |  |  |  |  |  |  |  |  |  |  |  |
| Exposure (e<sup>-</sup>/Å<sup>2</sup>) | 33 |  |  |  |  |  |  |  |  |  |  |  |  |  |
| Defocus range (µm) | -1.6 to -3.2 |  |  |  |  |  |  |  |  |  |  |  |  |  |
| Pixel size (Å) | 0.83 |  |  |  |  |  |  |  |  |  |  |  |  |  |
| Movies collected | 5,199 |  |  |  |  |  |  |  |  |  |  |  |  |  |
| Frames per movie | 20 |  |  |  |  |  |  |  |  |  |  |  |  |  |
| Data processing |  |  |  |  |  |  |  |  |  |  |  |  |  |  |
| Initial particles | 100,605 (C<sub>2</sub> symmetry-expanded: 201,210) |  |  |  |  |  |  |  |  |  |  |  |  |  |
| Final no. particles | 100,605 | 118,683 | 201,210 | 36,925 | 19,764 | 26,427 | 23,019 | 16,991 | 34,482 | 12,173 | 24,096 | 11,035 | 17,833 | 17,312 |
| Symmetry | C<sub>2</sub> | C<sub>1</sub> | C<sub>1</sub> | C<sub>2</sub> | C<sub>1</sub> | C<sub>1</sub> | C<sub>1</sub> | C<sub>1</sub> | C<sub>1</sub> | C<sub>1</sub> | C<sub>1</sub> | C<sub>1</sub> | C<sub>1</sub> | C<sub>1</sub> |
| Map resolution (Å) | 2.7 | 3.7 | 3.7 | 3.2 | 3.7 | 3.5 | 3.7 | 3.8 | 3.7 | 4.3 | 3.5 | 3.8 | 3.8 | 3.7 |
| Sharpening B factor | -46.2 | -74.4 | -92.5 | -49.8 | -61.8 | -61.1 | -57.6 | -45.6 | -58.0 | -73.8 | -54.5 | -65.2 | -54.9 | -61.7 |
| EMD ID |  |  |  |  |  |  |  |  |  |  |  |  |  |  |
| Model refinement statistics |  |  |  |  |  |  |  |  |  |  |  |  |  |  |
| CC (map/model) | 0.86 | 0.83 | 0.82 | 0.71 | 0.79 | 0.79 | 0.82 | 0.79 | 0.69 | 0.71 | 0.81 | 0.77 | 0.77 | 0.79 |
| Resolution (map/model) | 2.65 | 3.4 | 3.68 | 3.13 | 3.48 | 3.56 | 3.36 | 3.55 | 3.57 | 3.94 | 3.39 | 3.73 | 3.64 | 3.58 |
| No. of atoms | 76,690 | 19,669 | 12,083 | 251,552 | 129,568 | 129,568 | 129,568 | 129,568 | 129,568 | 129,563 | 129,563 | 129,563 | 129,563 | 129,566 |
| No. of residues | 4074 | 1285 | 767 | 15,356 | 7872 | 7872 | 7872 | 7872 | 7872 | 7872 | 7872 | 7872 | 7872 | 7872 |
| No. of lipids | 36 | 0 | 0 | 36 | 21 | 21 | 21 | 21 | 21 | 21 | 21 | 21 | 21 | 21 |
| No. of ATP/ADP | 0 | 0 | 0 | 10 | 5 | 5 | 5 | 5 | 5 | 5 | 5 | 5 | 5 | 5 |
| No. of Mg ions | 0 | 0 | 0 | 10 | 5 | 5 | 5 | 5 | 5 | 5 | 5 | 5 | 5 | 5 |
| B-factor (Å<sup>2</sup>) |  |  |  |  |  |  |  |  |  |  |  |  |  |  |
| - protein | 54.05 | 56.13 | 77.88 | 84.48 | 55.65 | 70.37 | 80.22 | 83.27 | 70.70 | 112.72 | 79.93 | 65.52 | 66.49 | 101.5 |
| - ligands | 50.57 | 58.25 | - | 69.94 | 40.99 | 72.29 | 63.18 | 78.43 | 63.76 | 75.25 | 74.47 | 61.79 | 46.55 | 83.68 |
| Rotamer outliers (%) | 0.44 | 0.40 | 0.31 | 0.22 | 0.42 | 0.09 | 0.18 | 0.26 | 0.58 | 0.18 | 0.27 | 0.48 | 0.42 | 0.39 |
| Ramachandran (%) |  |  |  |  |  |  |  |  |  |  |  |  |  |  |
| - outliers | 0.00 | 0.00 | 0.00 | 0.01 | 0.001 | 0.003 | 0.004 | 0.01 | 0.003 | 0.01 | 0.00 | 0.04 | 0.04 | 0.04 |
| - allowed | 1.57 | 1.91 | 1.59 | 1.56 | 1.52 | 1.65 | 1.44 | 1.49 | 1.49 | 1.67 | 1.58 | 1.47 | 1.65 | 1.79 |
| - favored | 98.43 | 98.08 | 98.41 | 98.42 | 98.47 | 98.34 | 98.56 | 98.49 | 98.48 | 98.31 | 98.42 | 98.49 | 98.31 | 98.17 |
| Clash score | 1.66 | 2.44 | 2.32 | 2.26 | 2.60 | 2.65 | 2.53 | 2.67 | 2.99 | 2.38 | 2.30 | 2.52 | 2.38 | 3.57 |
| MolProbity score | 0.92 | 1.03 | 1.01 | 1.00 | 1.05 | 1.05 | 1.04 | 1.05 | 1.09 | 1.02 | 1.01 | 1.04 | 1.02 | 1.15 |
| RMSD |  |  |  |  |  |  |  |  |  |  |  |  |  |  |
| - bonds (Å) | 0.004 | 0.004 | 0.02 | 0.003 | 0.003 | 0.003 | 0.004 | 0.003 | 0.003 | 0.002 | 0.003 | 0.003 | 0.003 | 0.003 |
| - angles (°) | 0.455 | 0.416 | 0.386 | 0.407 | 0.414 | 0.424 | 0.417 | 0.407 | 0.412 | 0.410 | 0.416 | 0.419 | 0.428 | 0.421 |
| EMRinger score | 5.11 | 3.96 | 1.61 | 2.56 | 3.24 | 2.95 | 3.32 | 2.85 | 3.32 | 1.35 | 2.89 | 2.32 | 2.49 | 2.8 |
| PDB ID |  |  |  |  |  |  |  |  |  |  |  |  |  |  |

**Extended Data Table 2.** Composition of *T. brucei* ATP synthase dimer.

| Subunit name | TriTrypDB Lister strain 427 ID | TriTrypDB TREU927 strain ID | Uniprot TREU927 strain ID | Residues | Residues built |
| --- | --- | --- | --- | --- | --- |
| **F<sub>1</sub> subcomplex** |  |  |  |  |  |
| $\alpha$ | Tb427_070081800<br>Tb427_070081900 | Tb927.7.7420<br>Tb927.7.7430 | Q57TX9 | 584 | 45-151,<br>161-584 |
| $\beta$ | Tb427_030013500 | Tb927.3.1380 | Q57XX1 | 519 | 26-514 |
| $\gamma$ | Tb427_100005200 | Tb927.10.180 | B0Z0F6 | 305 | 2-301 |
| $\delta$ | Tb427_060054900 | Tb927.6.4990 | Q586H1 | 182 | 22-182 |
| $\epsilon$ | Tb427_100054600 | Tb427.10.5050 | N/A | 75 | 11-75 |
| **p18** | Tb427_050022900 | Tb927.5.1710 | Q57ZP0 | 188 | 23-188 |
| **F<sub>0</sub> subcomplex** |  |  |  |  |  |
| **OSCP** | Tb427_100087100 | Tb927.10.8030 | Q38AG1 | 255 | 18-202,<br>208-255 |
| *a* | mt encoded | mt encoded | P24499 | 231 | 1-231 |
| *b* | Tb427_040009100 | Tb927.4.720 | Q580A0 | 105 | 26-105 |
| *c* | Tb427_100018700<br>Tb427_110057900<br>Tb427_070019000 | Tb927.10.1570<br>Tb927.11.5280<br>Tb927.7.1470 | Q38C84<br>Q385P0<br>Q57WQ3 | 118 | 41-118 |
| *d* | Tb427_050035800 | Tb927.5.2930 | Q57ZW9 | 370 | 17-325,<br>332-354 |
| *e* | Tb427_110010200 | Tb927.11.600 | N/A | 92 | 1-383 |
| *f* | Tb427_030016600 | Tb927.3.1690 | Q57ZE2 | 145 | 2-136 |
| *g* | Tb427_020016900 | Tb927.2.3610 | Q586X8 | 144 | 16-144 |
| *i/j* | Tb427_030029400 | Tb927.3.2880 | Q57ZM4 | 104 | 2-104 |
| *k* | Tb427_070011800 | Tb927.7.840 | Q57VT0 | 124 | 20-124 |
| **8** | Tb427_040037300 | Tb927.4.3450 | Q585K5 | 114 | 29-114 |
| **ATBTB1** | Tb427_100008400 | Tb927.10.520 | Q38CI8 | 396 | 1-383 |
| **ATPTB3** | Tb427_110067400 | Tb927.11.6250 | Q385E4 | 269 | 2-269 |
| **ATPTB4** | Tb427_100105100 | Tb927.10.9830 | Q389Z3 | 157 | 21-157 |
| **ATPTB6** | Tb427_110017200 | Tb927.11.1270 | Q387C5 | 169 | 2-169 |
| **ATPTB11** | Tb427_030021500 | Tb927.3.2180 | Q582T1 | 156 | 18-156 |
| **ATPTB12** | Tb427_050037400 | Tb927.5.3090 | Q57Z84 | 101 | 5-100 |
| **ATPEG3** | Tb427_060009300 | Tb927.6.590 | Q583U4 | 98 | 14-98 |
| **ATPEG4** | N/A | Tb927.11.2245 | N/A | 62 | 1-62 |

**Extended Data Table 3.**
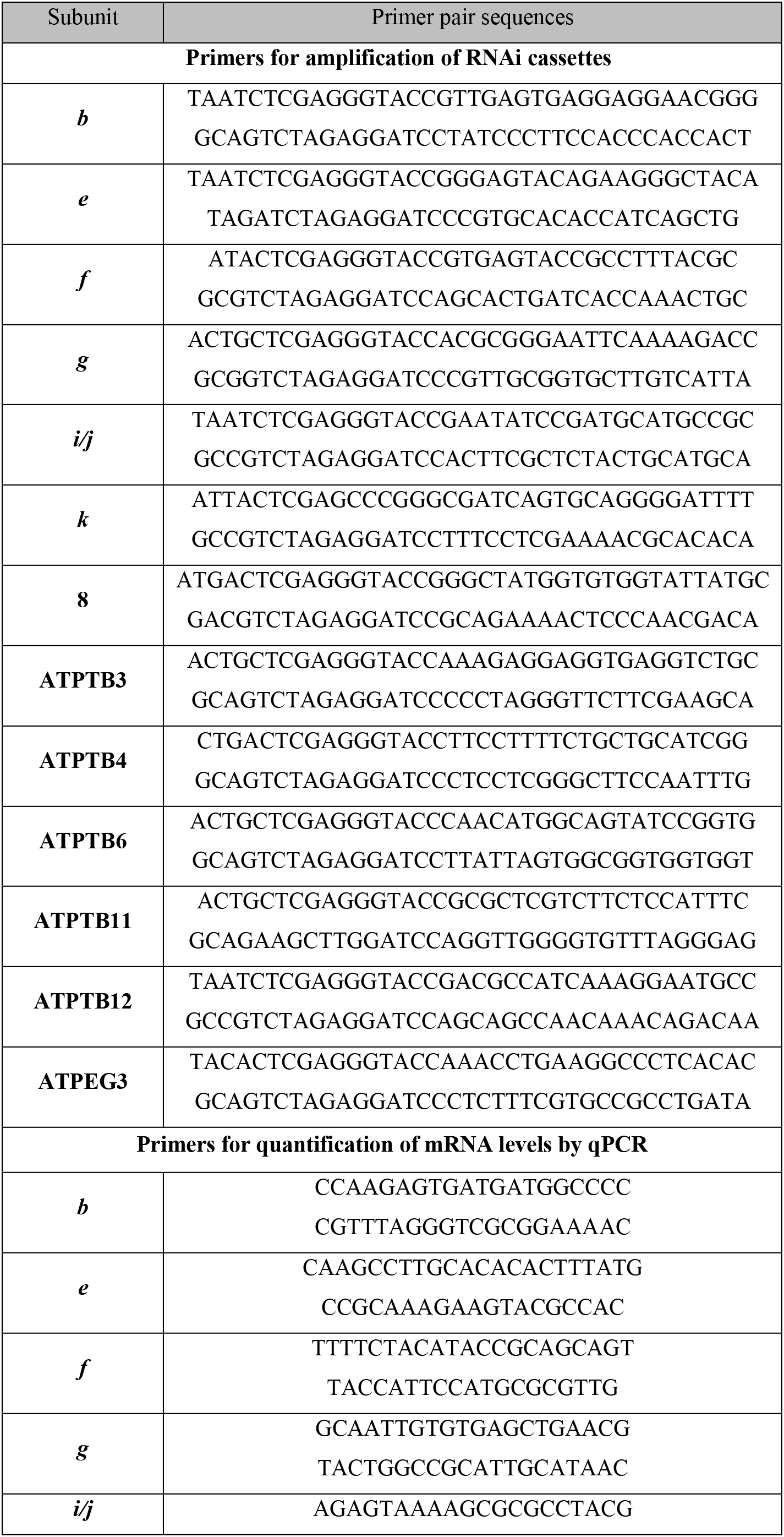
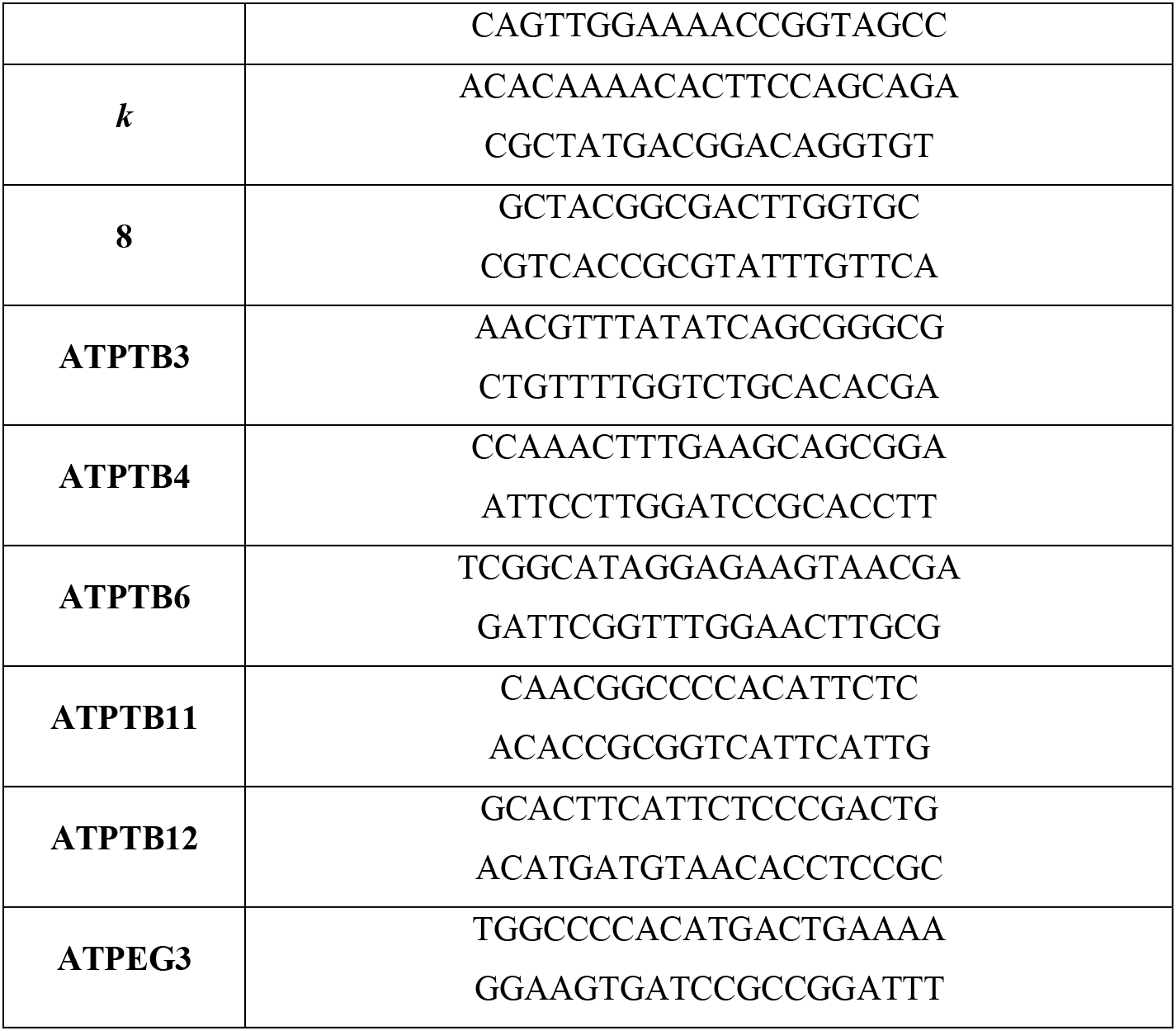
List of primers used in the study.

**Extended Data Table 4.** List of antibodies used in the study.

| Target | Type | Reference | Dilution SDS-PAGE | Dilution BN-PAGE |
| --- | --- | --- | --- | --- |
| **Primary antibodies** |  |  |  |  |
| **subunit-β** | rabbit polyclonal | 1 | 1:2000 | 1:2000 |
| **p18** | rabbit polyclonal | 1 | 1:1000 | - |
| **ATPTB1** | rabbit polyclonal | 1 | 1:1000 | 1:1000 |
| **subunit-*d*** | rabbit polyclonal | 1 | 1:1000 | 1:500 |
| **mtHsp70** | mouse monoclonal | 2 | 1:5000 | - |
| **Secondary antibodies** |  |  |  |  |
| **goat anti-rabbit IgG HRP conjugate** |  | BioRad 1721019 | 1:2000 | 1:2000 |
| **goat anti-mouse IgG HRP conjugate** |  | BioRad 1721011 | 1:2000 | 1:2000 |

## References

1. Paumard, P. et al. The ATP synthase is involved in generating mitochondrial cristae morphology. EMBO J 21, 221–30 (2002).

2. Davies, K.M., Anselmi, C., Wittig, I., Faraldo-Gomez, J.D. & Kuhlbrandt, W. Structure of the yeast F_1_F_o_-ATP synthase dimer and its role in shaping the mitochondrial cristae. Proc Natl Acad Sci U S A 109, 13602–7 (2012).

3. Panek, T., Elias, M., Vancova, M., Lukes, J. & Hashimi, H. Returning to the Fold for Lessons in Mitochondrial Crista Diversity and Evolution. Curr Biol 30, R575–R588 (2020).

4. Kuhlbrandt, W. Structure and Mechanisms of F-Type ATP Synthases. Annu Rev Biochem 88, 515–549 (2019).

5. Spikes, T.E., Montgomery, M.G. & Walker, J.E. Structure of the dimeric ATP synthase from bovine mitochondria. Proc Natl Acad Sci U S A 117, 23519–23526 (2020).

6. Pinke, G., Zhou, L. & Sazanov, L.A. Cryo-EM structure of the entire mammalian F-type ATP synthase. Nat Struct Mol Biol 27, 1077–1085 (2020).

7. Guo, H., Bueler, S.A. & Rubinstein, J.L. Atomic model for the dimeric F_o_ region of mitochondrial ATP synthase. Science 358, 936–940 (2017).

8. Murphy, B.J. et al. Rotary substates of mitochondrial ATP synthase reveal the basis of flexible F_1_-F_o_ coupling. Science 364, eaaw9128 (2019).

9. Flygaard, R.K., Mühleip, A., Tobiasson, V. & Amunts, A. Type III ATP synthase is a symmetry-deviated dimer that induces membrane curvature through tetramerization. Nature Communications 11, 5342 (2020).

10. Muhleip, A., McComas, S.E. & Amunts, A. Structure of a mitochondrial ATP synthase with bound native cardiolipin. Elife 8, e51179 (2019).

11. Mühleip, A. et al. ATP synthase hexamer assemblies shape cristae of *Toxoplasma* mitochondria. Nature Communications 12, 120 (2021).

12. Gahura, O. et al. The F_1_-ATPase from *Trypanosoma brucei* is elaborated by three copies of an additional p18-subunit. FEBS J 285, 614–628 (2018).

13. Montgomery, M.G., Gahura, O., Leslie, A.G.W., Zikova, A. & Walker, J.E. ATP synthase from *Trypanosoma brucei* has an elaborated canonical F_1_-domain and conventional catalytic sites. Proc Natl Acad Sci U S A 115, 2102–2107 (2018).

14. Serricchio, M. et al. Depletion of cardiolipin induces major changes in energy metabolism in *Trypanosoma brucei* bloodstream forms. FASEB J 35, 21176 (2020).

15. Muhleip, A.W., Dewar, C.E., Schnaufer, A., Kuhlbrandt, W. & Davies, K.M. In situ structure of trypanosomal ATP synthase dimer reveals a unique arrangement of catalytic subunits. Proc Natl Acad Sci U S A 114, 992–997 (2017).

16. Schnaufer, A., Clark-Walker, G.D., Steinberg, A.G. & Stuart, K. The F_1_-ATP synthase complex in bloodstream stage trypanosomes has an unusual and essential function. EMBO J 24, 4029–40 (2005).

17. Brown, S.V., Hosking, P., Li, J. & Williams, N. ATP synthase is responsible for maintaining mitochondrial membrane potential in bloodstream form *Trypanosoma brucei*. Eukaryot Cell 5, 45–53 (2006).

18. Gahura, O., Hierro-Yap, C. & Zikova, A. Redesigned and reversed: Architectural and functional oddities of the trypanosomal ATP synthase. Parasitology 148, 1151–1160 (2021).

19. Hierro-Yap, C. et al. Bioenergetic consequences of F_o_F_1_-ATP synthase/ATPase deficiency in two life cycle stages of *Trypanosoma brucei*. J Biol Chem 296, 100357 (2021).

20. Gahura, O., Panicucci, B., Vachova, H., Walker, J.E. & Zikova, A. Inhibition of F_1_-ATPase from *Trypanosoma brucei* by its regulatory protein inhibitor TbIF_1_. FEBS J 285, 4413–4423 (2018).

21. Zikova, A., Schnaufer, A., Dalley, R.A., Panigrahi, A.K. & Stuart, K.D. The F(0)F(1)-ATP synthase complex contains novel subunits and is essential for procyclic *Trypanosoma brucei*. PLoS Pathog 5, e1000436 (2009).

22. Perez, E. et al. The mitochondrial respiratory chain of the secondary green alga *Euglena gracilis* shares many additional subunits with parasitic Trypanosomatidae. Mitochondrion 19 Pt B, 338–49 (2014).

23. Sathish Yadav, K.N. et al. Atypical composition and structure of the mitochondrial dimeric ATP synthase from *Euglena gracilis*. Biochim Biophys Acta 1858, 267–275 (2017).

24. Dewar, C.E., Oeljeklaus, S., Wenger, C., Warscheid, B. and Schneider, A. Characterisation of a highly diverged mitochondrial ATP synthase F_o_ subunit in *Trypanosoma brucei*. J Biol Chem. Epub ahead of print, 101829 (2022)

25. Aphasizheva, I. et al. Lexis and Grammar of Mitochondrial RNA Processing in Trypanosomes. Trends Parasitol 36, 337–355 (2020).

26. Blum, B., Bakalara, N. & Simpson, L. A model for RNA editing in kinetoplastid mitochondria: “guide” RNA molecules transcribed from maxicircle DNA provide the edited information. Cell 60, 189–98 (1990).

27. Adler, B.K., Harris, M.E., Bertrand, K.I. & Hajduk, S.L. Modification of *Trypanosoma brucei* mitochondrial rRNA by posttranscriptional 3’ polyuridine tail formation. Mol Cell Biol 11, 5878–84 (1991).

28. Hofer, A., Steverding, D., Chabes, A., Brun, R. & Thelander, L. *Trypanosoma brucei* CTP synthetase: a target for the treatment of African sleeping sickness. Proc Natl Acad Sci U S A 98, 6412–6 (2001).

29. Sobti, M., Walshe, J.L., Wu, D. et al. Cryo-EM structures provide insight into how *E. coli* F_1_F_o_ ATP synthase accommodates symmetry mismatch. Nat Commun 11, 2615 (2020). https://doi.org/10.1038/s41467-020-16387-2

30. Gupta, K. et al. The role of interfacial lipids in stabilizing membrane protein oligomers. Nature 541, 421–424 (2017).

31. Arnold, I., Pfeiffer, K., Neupert, W., Stuart, R.A. & Schagger, H. Yeast mitochondrial F_1_F_o_-ATP synthase exists as a dimer: identification of three dimer-specific subunits. EMBO J 17, 7170–8 (1998).

32. Gu, J. et al. Cryo-EM structure of the mammalian ATP synthase tetramer bound with inhibitory protein IF_1_. Science 364, 1068–1075 (2019).

33. Spikes, T.E., Montgomery, M.G. & Walker, J.E. Interface mobility between monomers in dimeric bovine ATP synthase participates in the ultrastructure of inner mitochondrial membranes. Proc Natl Acad Sci U S A 118, e2021012118 (2021).

34. Cadena, L.R. et al. Mitochondrial contact site and cristae organization system and F_1_Fo-ATP synthase crosstalk is a fundamental property of mitochondrial cristae. mSphere 6, e0032721 (2021).

35. Davies, K.M. et al. Macromolecular organization of ATP synthase and complex I in whole mitochondria. Proc Natl Acad Sci U S A 108, 14121–6 (2011).

36. Blum, T.B., Hahn, A., Meier, T., Davies, K.M. & Kühlbrandt, W. Dimers of mitochondrial ATP synthase induce membrane curvature and self-assemble into rows. Proc Natl Acad Sci U S A 116, 4250–4255 (2019).

37. Bochud-Allemann, N. & Schneider, A. Mitochondrial substrate level phosphorylation is essential for growth of procyclic *Trypanosoma brucei*. J Biol Chem 277, 32849–54 (2002).

38. Poon, S.K., Peacock, L., Gibson, W., Gull, K. & Kelly, S. A modular and optimized single marker system for generating *Trypanosoma brucei* cell lines expressing T7 RNA polymerase and the tetracycline repressor. Open Biol 2, 110037 (2012).

39. Allemann, N. & Schneider, A. ATP production in isolated mitochondria of procyclic Trypanosoma brucei. Mol Biochem Parasitol 111, 87–94 (2000).

40. Aibara, S., Dienemann, C., & Cramer, P.. Structure of an inactive RNA polymerase II dimer. Nucleic Acids Research, gkab783 (2021).

41. de la Rosa-Trevin, J.M. et al. Scipion: A software framework toward integration, reproducibility and validation in 3D electron microscopy. J Struct Biol 195, 93–9 (2016).

42. Zhang, K. Gctf: Real-time CTF determination and correction. J Struct Biol 193, 1–12 (2016).

43. Cowtan, K. The Buccaneer software for automated model building. 1. Tracing protein chains. Acta Crystallogr D Biol Crystallogr 62, 1002–11 (2006).

44. Waterhouse, A. et al. SWISS-MODEL: homology modelling of protein structures and complexes. Nucleic Acids Res 46, W296–W303 (2018).

45. Williams, C.J., Headd, J.J., Moriarty, N.W., Prisant, M.G., Videau, L.L., Deis, L.N., Verma, V., Keedy, D.A., Hintze, B.J., Chen, V.B. and Jain, S. MolProbity: More and better reference data for improved all-atom structure validation. Protein Science, 27, 293–315 (2018).

46. Barad, B.A., Echols, N., Wang, R.Y.R., Cheng, Y., DiMaio, F., Adams, P.D. and Fraser, J.S. EMRinger: side chain–directed model and map validation for 3D cryo-electron microscopy. Nature methods, 12, 943–946 (2015).

47. Goddard, T.D. et al. UCSF ChimeraX: Meeting modern challenges in visualization and analysis. Protein Sci 27, 14–25 (2018).

48. Ho, B.K. & Gruswitz, F. HOLLOW: generating accurate representations of channel and interior surfaces in molecular structures. BMC Struct Biol 8, 49 (2008).

## Extended Data references

1. Muhleip, A., McComas, S.E. & Amunts, A. Structure of a mitochondrial ATP synthase with bound native cardiolipin. Elife 8, e51179 (2019).

2. Larkin, M.A. et al. (2007). Clustal W and Clustal X version 2.0. Bioinformatics, 23, 2947–2948 (2007).

3. Burki, F., Roger, A.J., Brown, M.W. & Simpson, A.G.B. The New Tree of Eukaryotes. Trends Ecol Evol 35, 43–55 (2020).

4. Protein Sequence Similarity Search. Curr Protoc Bioinformatics 60, 3151–31523 (2017).

5. Huang, Y., Niu, B., Gao, Y., Fu, L. & Li, W. CD-HIT Suite: a web server for clustering and comparing biological sequences. Bioinformatics 26, 680–2 (2010).

6. Sievers, F. et al. Fast, scalable generation of high-quality protein multiple sequence alignments using Clustal Omega. Mol Syst Biol 7, 539 (2011).

7. Edgar, R.C. MUSCLE: multiple sequence alignment with high accuracy and high throughput. Nucleic Acids Res 32, 1792–7 (2004).

